# Developmental increase of inhibition drives decorrelation of neural activity

**DOI:** 10.1101/2021.07.06.451299

**Authors:** Mattia Chini, Thomas Pfeffer, Ileana L. Hanganu-Opatz

**Author notes:** Corresponding author: Mattia Chini.

## Abstract

Throughout development, the brain transits from early highly synchronous activity patterns to a mature state with sparse and decorrelated neural activity, yet the mechanisms underlying this process are unknown. The developmental transition has important functional consequences, as the latter state allows for more efficient storage, retrieval and processing of information. Here, we show that, in the mouse medial prefrontal cortex (mPFC), neural activity during the first two postnatal weeks decorrelates following specific spatial patterns. This process is accompanied by a concomitant tilting of excitation/inhibition (E-I) ratio towards inhibition. Using optogenetic manipulations and neural network modeling, we show that the two phenomena are mechanistically linked, and that a relative increase of inhibition drives the decorrelation of neural activity. Accordingly, in two mouse models of neurodevelopmental disorders, subtle alterations in E-I ratio are associated with specific impairments in the correlational structure of spike trains. Finally, capitalizing on EEG data from newborn babies, we show that an analogous developmental transition takes place also in the human brain. Thus, changes in E-I ratio control the (de)correlation of neural activity and, by these means, its developmental imbalance might contribute to the pathogenesis of neurodevelopmental disorders.

## Introduction

Neural activity in the developing brain has several unique traits such as discontinuity^1^, extremely low firing rates^2^, a loose temporal coordination of excitation and inhibition^3^, and weak modulation by behavioral state^4,5^. These patterns of early activity have been described in humans^6^ as well as in disparate model organisms ranging from fish^7^ to flies^8^, and from rodents^9^ to brain organoids^10^. As brain networks mature, they gradually evolve into exhibiting motives with adult-like spatiotemporal properties. Oscillatory events become more rhythmic, increasing their amplitude and average frequency^11^, oscillation patterns become more complex^10^, the ratio of excitatory and inhibitory conductances (E-I ratio) decreases^12^, excitation and inhibition tighten on a temporal scale^3,13^, and brain activity decorrelates and sparsifies^14,15^. The relationship between decorrelation and E-I ratio has been the subject of extensive experimental and theoretical work in the adult brain^16–18^. Decorrelated and sparse activity is a hallmark of adult spike trains^19,20^ and artificial neural networks alike^21–23^. This activity pattern bears important functional and behavioral relevance, as it allows for efficient storing and retrieval of information, while minimizing energy consumption^19,21,22^. However, it is still unknown whether changes in E-I ratio underlie the developmental decorrelation of brain activity. The (patho)physiological relevance of this process is underscored by the hypothesis that altered E-I ratio is the hallmark of neurodevelopmental disorders, such as autism^24–28^ and schizophrenia^27,29^, diseases that have also been linked to disruption of correlated activity in animal models^30–32^.

E-I ratio is controlled by the interplay between pyramidal neurons (PYRs) and interneurons (INs). Throughout development, both populations of neurons migrate into the cortex following an “inside-out” sequence that corresponds to their birthdate^33,34^. This process is guided by cues provided by PYRs, that populate the cortical layers at an earlier time point^33^. The functional integration of INs into the cortical circuitry is a slow process that is initiated by the establishment of transient circuits, mainly including somatostatin-positive (SST^+^) INs^35,36^. At later time points, parvalbumin-positive (PV^+^) INs are also integrated into local networks^35–37^. In rodents, the development of inhibitory synapses is not complete until postnatal day (P)30^38^. It is thus conceivable that the developmental strengthening of inhibitory synapses and the ensuing tilting of E-I ratio towards inhibition^12^ might underlie the decorrelation of neural activity. In favor of this hypothesis, chronic manipulation of IN activity in the murine barrel cortex resulted in altered temporal and spatial structure of brain activity^39–41^. However, direct evidence linking E-I ratio and the strengthening of inhibition with the developmental decorrelation of brain activity is still lacking.

Here, we combined in vivo extracellular electrophysiological recordings and optogenetics with neural network modeling to systematically explore the relationship between E-I ratio and the decorrelation of neural activity in the developing rodent and human brain. In the murine mPFC, a brain area where E-I ratio is of particular relevance in the context of neurodevelopmental disorders^24^, we show that inhibition increases and spike trains decorrelate throughout early development. Using neural network modeling and bidirectional optogenetic manipulation of IN activity, we further uncover how inhibition increase drives the change in the correlation structure. Moreover, in two mouse models of impaired neurodevelopment, we report that excessively decreased developmental E-I ratio results in lower spike train correlations. Finally, we investigate two different EEG datasets and illustrate the translational relevance of these findings by providing first insights into analogous developmental processes that take place in newborn babies.

## Results

### The patterns of prefrontal activity dynamically evolve with age

To investigate the relationship between E-I ratio and the decorrelation of neural activity that occurs throughout development, we interrogated a large dataset (n=117 mice) of multi-site extracellular recordings of local field potential (LFP) and single activity (SUA) from the prelimbic subdivision of the mPFC of unanesthetized P2-P12 mice (Figure 1A). Across this developmental phase, the LFP evolves from an almost complete lack of activity (silent periods) to uninterrupted (continuous) activity, passing through intermediate stages in which silent periods alternate with bouts of neuronal activity (active periods) (Figure 1A). To quantify this transition, we calculated the proportion of active periods over the recording and found that it monotonically increases over age (age slope=0.88, 95% C.I. [0.84; 0.93], p<10^-50^, generalized linear model) (Figure 1B). The increase in the proportion of active periods resulted from the augmentation of both number and duration of individual active periods until continuous activity was detected (Supp. Figure 1A-C). Accompanying these high-level changes in activity dynamics, the maximum amplitude of active periods, the broadband LFP power, and the SUA firing rate exponentially increased over age (age slope=0.24, 0.47 and 0.21, 95% C.I. [0.18; 0.30] [0.40; 0.53] and [0.15; 0.27], p<10^-9^, p<10^-23^, p<10^-11^, respectively, generalized linear model) (Figure 1C, Supp. Figure 1D-F).

**Figure 1.**
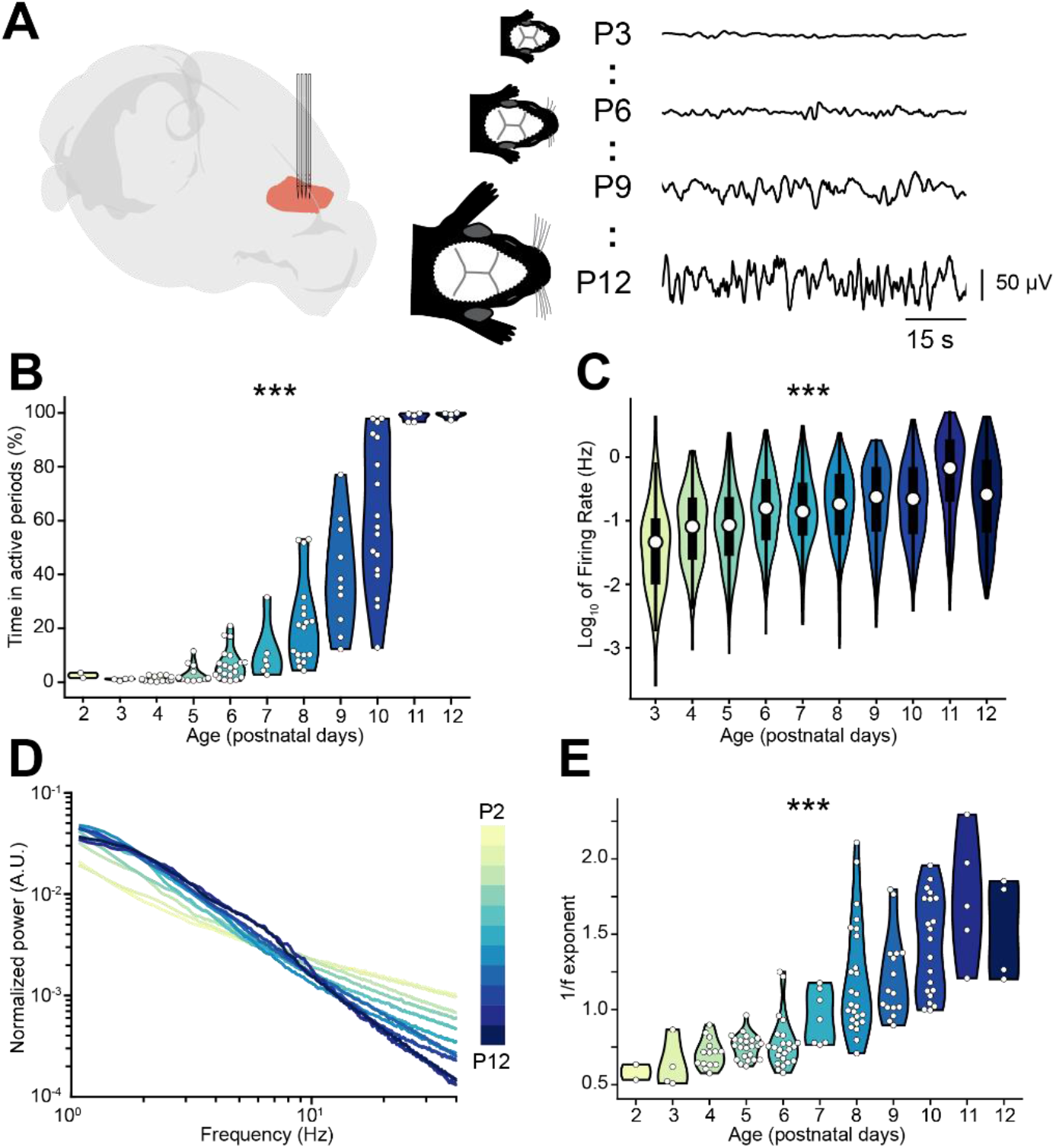
Active periods and LFP properties of the mouse mPFC across the first two postnatal weeks. (**A**) Schematic representation^42^ of extracellular recordings in the mPFC of P2-12 mice (left), and representative LFP traces from P3, P6, P9 and P12 mice (right). (**B** and **C**) Violin plots displaying the percentage of time spent in active periods (B) and the SUA firing rate (C) of P2-12 mice (n=117 mice and 2269 single units, respectively). (**D**) Log-log plot displaying the normalized median PSD power in the 1-40 Hz frequency range of P2-12 mice (n=117 mice). Color codes for age with one day increment. (**E**) Violin plot displaying the 1/f exponent of P2-12 mice (n=117 mice). In (B) and (E) white dots indicate individual data points. In (C) data are presented as median, 25^th^, 75^th^ percentile and interquartile range. In (B), (C) and (E) the shaded area represents the probability distribution density of the variable. In (D) data are presented as median. Asterisks in (B), (C) and (E) indicate significant effect of age. *** p < 0.001. Generalized linear models (B-C) and linear model (E). For detailed statistical results, see S1 Table.

Changes in the log-log power spectral density (PSD) slope (reflected by the 1/f exponent) have been linked to E-I ratio by several experimental^24,43,44^ and theoretical studies^24,45,46^. In particular, a relative increase in inhibition is thought of leading to a steeper PSD slope (higher 1/f exponent), whereas the opposite occurs when E-I ratio shifts towards excitation. Given that INs are thought to have a more protracted integration into cortical circuits than PYRs, we reasoned that this might be accompanied by a developmental shift of the E-I ratio towards inhibition. In line with this hypothesis, the PSD slope grew steeper over age, as readily observed when the area under the curve of the PSD was normalized (Figure 1D). To quantify this observation, we parameterized the PSDs using a recently published protocol^47^, and confirmed that the 1/f exponent increases over age (age slope=0.12, 95% C.I. [0.11; 0.14], p<10^-27^, linear model) (Figure 1E).

These data monitor the age-dependent dynamics of LFP and SUA in the mouse mPFC and lead to the hypothesis that, throughout development, E-I ratio tilts towards inhibition.

### E-I ratio controls pairwise spike train correlations in a neural network model

To explore the relationship between the 1/f exponent, E-I ratio and the (de)correlation of neuronal spike trains, we simulated a neural network of 400 interconnected conductance-based leaky integrate-and-fire (LIF) neurons (Figure 2A). In line with anatomical data^48,49^, 80% of those simulated neurons were excitatory (PYRs), whereas 20% were inhibitory (INs). PYRs were simulated with outgoing excitatory AMPA synapses, while INs were simulated with outgoing inhibitory GABAergic synapses, including recurrent connections for both PYRs and INs. Both neuron types received input noise and PYRs received an additional external excitatory Poisson stimulus with a constant spike rate of 1.5 spikes / second (see Methods for details on the model). We parametrically varied the AMPA and GABA conductances on both PYRs and INs and defined the network’s net inhibition strength as the ratio between the inhibitory and excitatory conductances. The network was simulated for 50 seconds for each parameter combination. The network’s LFP was defined as the sum of the absolute values of all synaptic currents on PYRs, which was shown to be a reliable proxy of experimental LFP recordings^24,50^. Across all parameters combinations, INs exhibited higher average firing rates compared to PYRs (IN: 2.87 Hz, StD = 0.17 Hz; PYR: 1.76 Hz, StD = 0.099 Hz).

**Figure 2.**
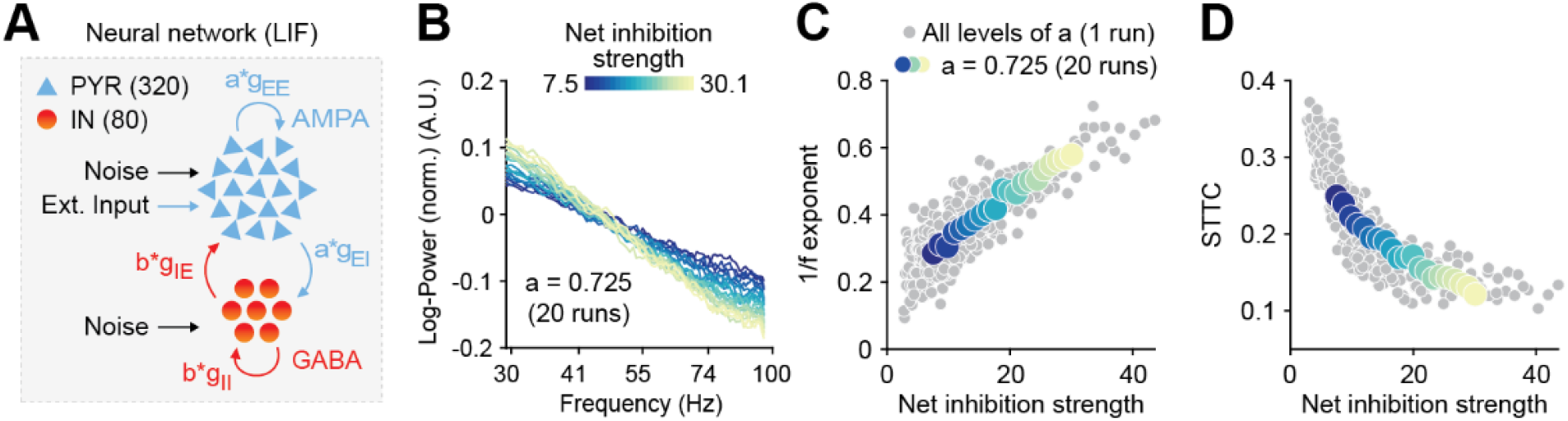
Increased inhibition leads to an increase in the 1/f exponent and decorrelates spike trains in a neural network model. (**A**) Schematic representation of the neural network model. (**B**) Log-log plot displaying the normalized median PSD power in the 30-100 Hz frequency range for varying level of inhibition. Color codes for inhibition strength. (**C**) Scatter plot displaying the 1/f exponent as a function of net inhibition strength. (**D**) Scatter plot displaying average STTC as a function of net inhibition strength. For (C) and (D) color codes for inhibition strength with fixed excitation level.

In agreement with previous results^24,46^, increases in the network’s net inhibition strength robustly tilted the PSD slope (in the range from 30 – 100 Hz). Increased levels of net inhibition were associated with a steeper decay of power as a function of frequency (Figure 2B) and a corresponding increase of the 1/f exponent, across a range of AMPA conductance levels (Figure 2C; Person correlation coefficient, averaged across AMPA levels: r = 0.768; 95% C.I. [0.50; 0.92]). We next examined the effect of increasing the network’s net inhibition strength on neural correlations. To this end, we computed the spike time tiling coefficient (STTC; at a lag of 1 s), a parameter that measures pairwise correlations between spike trains without being biased by firing rate^51^, on the network’s spike matrices (both PYRs and INs) and across all levels of net inhibition strength. For all levels of AMPA conductance, we found that increased net inhibition strength results in a robust decrease in STTC (Figure 2D; Person correlation coefficient, averaged across AMPA levels: r = −0.76; 95% C.I. [−0.89; −0.63]).

Thus, simulations of a biologically plausible neural circuit reveal that increased net inhibition strength leads to an increase of the PSD 1/f exponent that is accompanied by decorrelation of neural spike trains.

### Prefrontal spike trains decorrelate over development

Since neural network modeling predicts that a shift of E-I ratio towards inhibition leads to higher 1/f exponent and decorrelation of neural activity, we tested on the experimental data whether the developmental increase in the 1/f exponent in the mouse mPFC was accompanied by a decorrelation of neural activity. For this, we calculated the STTC between >40.000 pairs of spike trains over a large range of lags (2.5 ms to 10 s) (Figure 3A). For the analysis, we only considered SUA that was recorded for at least60 min. To verify the robustness of STTC as an estimator, we compared the STTC obtained on the first and the second half of the recording. The STTCs computed on the two halves of the recording strongly correlated with each other across all the investigated lags (0.70, [0.50; 0.80] median and min-max Pearson correlation; 0.70 [0.52; 0.79] median and min-max Spearman correlation) (Figure 3B, Supp. Figure 2A), thus corroborating its robustness as an estimator. Throughout the manuscript, we will consider STTC computed at 1 s, yet the summary plots and the supplementary statistical table include values calculated at all lags.

**Figure 3.**
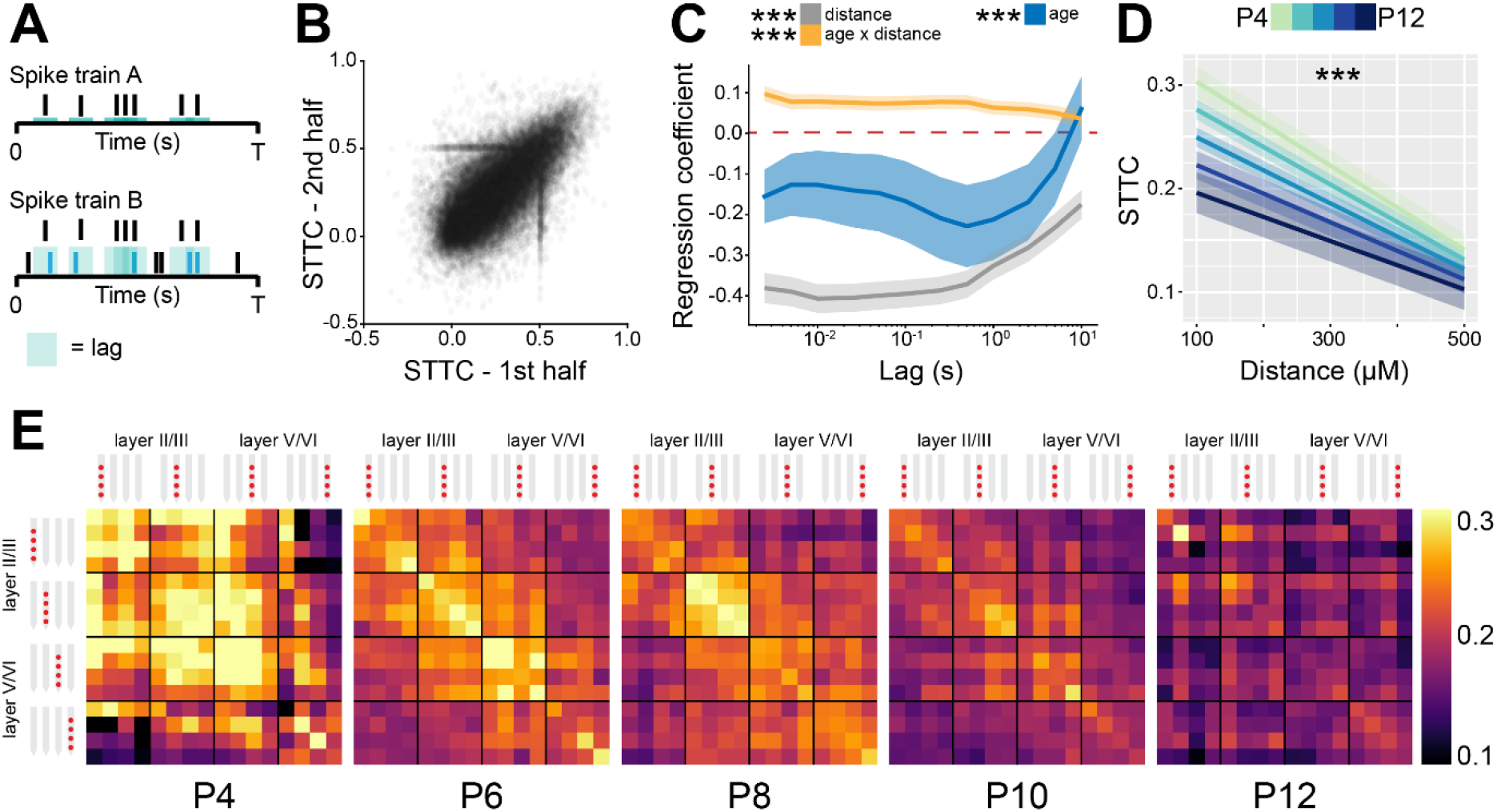
STTC decreases throughout development with a specific spatial profile in the mouse mPFC. (**A**) Schematic representation of the STTC quantification. (**B**) Scatter plot displaying the STTC computed in the 1^st^ half of the recording with respect to STTC computed in the 2^nd^ half of the recording (n=40921 spike train pairs and 82 mice). (**C**) Multivariate linear regression coefficients with respect to STTC lag (n=40921 spike train pairs and 82 mice). (**D**) Average STTC at 1 s lag of P4, P6, P8, P10 and P12 mice over distance (n=40921 spike train pairs and 82 mice). Color codes for age. (**E**) Weighted adjacency matrices displaying average STTC at 1 s lag of P4, P6, P8, P10 and P12 mice as a function of the recording sites in which the spike train pair has been recorded. Color codes for STTC value. In (C) regression coefficients are presented as mean and 95% C.I. In (D) data are presented as mean ± SEM. Asterisks in (C) indicate significant regression coefficients of the respective (interaction between) variables for STTC at 1 s lag. Asterisks in (D) indicate significant effect of age*distance interaction. *** p < 0.001. Linear mixed-effect models. For detailed statistical results, see S1 Table.

Using multivariate mixed hierarchical linear regression, we found that STTC negatively correlated with the distance between neurons (i.e. nearby neurons had higher STTC values than neurons that are far apart) over all the investigated lags (main distance effect, p<10^-78^ at 1 s lag, linear mixed-effect model) (Figure 3C-E). This is in line with previous studies conducted in in the adult^52–54^ and developing brain^14,51,55^ of several mammalian species. Further, STTC values negatively correlated with age at lags ≤ 5 s (main age effect. p<10^-4^ at 1 s lag, linear mixed-effect model), an effect that was strongest in the 100-1000 ms range (Figure 3C-E). This developmental STTC decrease did not occur uniformly across all neuron pairs. Rather, age and distance had a significant interaction, nearby pairs of neurons displaying a more severe decorrelation over age than neurons that were further apart (age*distance interaction, p<10^-12^ at 1 s lag, linear mixed-effect model) (Figure 3C-E).

Next, we fitted a separate statistical model that considers the “spatial configuration” of the neuron pairs (Supp. Figure 2B, see Materials and Methods for details). Even after accounting for the distance between neuron pairs, particularly at short lags, we found that “local” (i.e. from the same recording site) and “lateral” (i.e. from the same layer) spike train pairs displayed the highest STTC values (STTC=0.73 and 0.70, 95% C.I. [0.66; 0.79] and [0.64; 0.76], respectively, linear mixed-effect model). “Translaminar” pairs (i.e. from the same “column”) and pairs that did not fall in any of the previous categories (“other”) had the lowest STTC values (STTC=0.61 and 0.61, 95% C.I. [0.55; 0.67] and [0.55; 0.67], respectively, linear mixed-effect model) (Supp. Figure 2C-E). Further, “local” and “lateral” neuron pairs had a stronger developmental decrease in STTC values than “translaminar” and “other” pairs, particularly at short lags (Supp. Figure 2F, see S1 Table for details).

Taken together, these data indicate that, throughout development, as E-I ratio tilts towards inhibition, there is a concomitant decorrelation of pairwise neuronal activity computed over lags that span more than three orders of magnitude. This result is in agreement with data from the rodent barrel cortex^14,15^. In addition, we report that this process follows a specific spatial and structural pattern, with the activity of nearby neurons that are in the same cortical layer being the most affected.

### Optogenetic interneuron manipulation confirms developmental increase of inhibition

To experimentally substantiate the experimental evidence supporting the developmental increase of inhibition in the mouse mPFC, we optogenetically manipulated IN activity at different stages of early development. To this aim, we selectively transfected Dlx5/6^+^ and Gad2^+^ INs with either an excitatory (ChR2, n=19 mice) or an inhibitory opsin (ArchT, n=40 mice) using a combination of mouse lines and viral approaches. Briefly, expression of an excitatory opsin in INs was achieved by injecting P0-1 Dlx5/6^+^ and Gad2^+^ mice with a virus encoding for ChR2 (AAV9-Ef1alpha-DIO-hChR2(ET/TC)-eYFP). Expression of an inhibitory opsin was instead achieved by crossing Dlx5/6^+^ and Gad2^+^ mice with a mouse line (Ai40(RCL-ArchT/EGFP)-D) expressing ArchT under a cre-dependent promoter. No significant differences between experiments targeting Dlx5/6^+^ and Gad2^+^ neurons were detected and therefore, the datasets were pooled. In line with previously developed protocols^11,56,57^, we applied a 3 second-long “ramp-like” optogenetic stimulation of increasing intensity. This stimulation pattern gradually modulated the firing of the population of interest, while leading to minimal LFP artefacts^58,59^(Figure 4A, Supp. Figure 3A).

**Figure 4.**
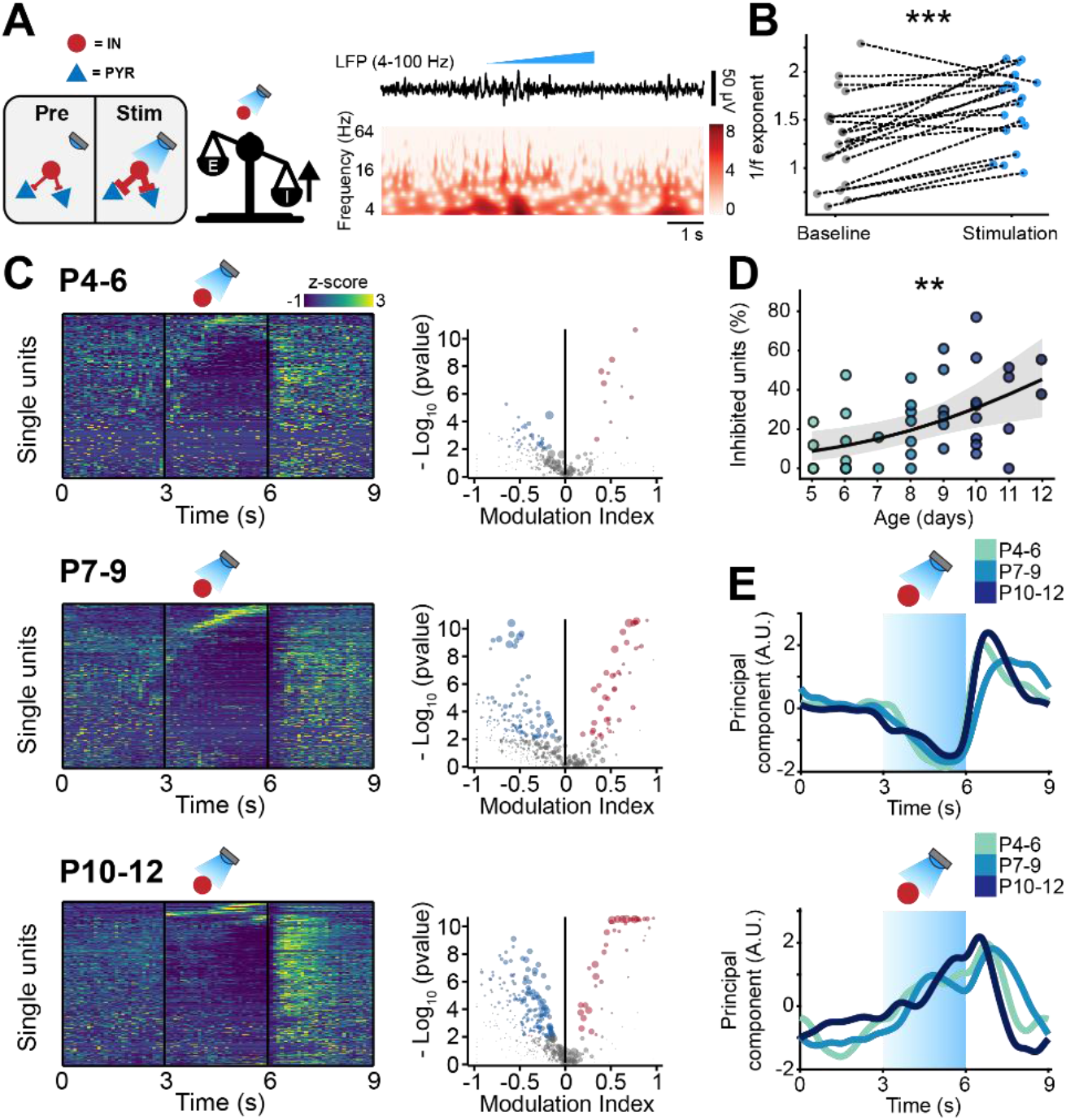
Optogenetic stimulation of IN activity leads to widespread inhibition in the developing mouse mPFC. (**A**) Schematic representation of the effects induced by optogenetic IN stimulation (left). Representative LFP trace (4-100 Hz band-pass filter) with a corresponding wavelet spectrum at an identical timescale during ramp light stimulation (473 nm, 3 s) of INs in the mPFC of a P10 mouse. (**B**) Scatter plot displaying the 1/f exponent during baseline and IN optogenetic stimulation (n=19 mice). (**C**) Z-scored single unit firing rates in response to optogenetic stimulation of INs (left) and volcano plot displaying the modulation index of pre vs stim single unit firing rates (right) for P4-6 (top, n=268 units and 5 mice), P7-9 (middle, n=480 units and 7 mice) and P10-12 (bottom, n=475 units and 7 mice) mice. Color codes for firing rate. (**D**) Scatter plot displaying the percentage of inhibited units with respect to age (n=19 mice). (**E**) 1^st^ (top, putative PYRs) and 2^nd^ (bottom, putative INs) PCA component of trial-averaged spike trains in response to optogenetic stimulation of INs. Color codes for age group. In (D) the regression is presented as mean and 95% C.I. Asterisks in (B) and (D) indicate significant effect of IN activation and age, respectively. ** p < 0.01, *** p < 0.001. Linear model (B) and generalized linear mixed-effect model (D). For detailed statistical results, see S1 Table.

While the LFP power was only mildly affected upon IN stimulation (Supp. Figure 3B), activation of INs increased the 1/f exponent, indicative of a shift in E-I ratio towards inhibition (main IN stimulation effect, p<10^-3^, linear model) (Figure 4B). In contrast to the minor effects on LFP, IN activation led to conspicuous modulation of SUA activity. Upon stimulus, a small number of neurons (putative INs) gradually increased their firing rate (Figure 4C). The proportion of stimulated INs was similar among mouse lines (main mouse line effect, p=0.14, generalized linear mixed-effect model) and across ages (main age effect=0.14, 95% C.I. [−0.04; 0.34], p=0.12, generalized linear mixed-effect model) (Supp. Figure 3C). This is in line with the histological quantification of the number of virally-transfected neurons that led to similar results for all mouse lines (main mouse line effect, p=0.45, linear mixed-effect model) and developmental stages (main age effect=-1.35, 95% C.I. [−3.22; 0.51], p=0.18, linear mixed-effect model) (Supp. Figure 3D). While putative INs increased their firing rate in response to optogenetic stimulation, a larger proportion of neurons (putative PYRs) significantly decreased their firing rate (Figure 4B). In line with the results above that indicated increasing inhibition throughout development, the proportion of inhibited neurons augmented with age (main age effect=0.31, 95% C.I. [0.09; 0.55], p=0.005, generalized linear mixed-effect model) (Figure 4D). Regardless of age, after terminating the optogenetic stimulus, PYRs responded with a prominent “rebound” increase in firing rate, similar to the effects reported for the adult brain^60,61^.

To dissect the main “neuronal trajectories” in response to light stimulation, we performed principal component analysis (PCA) on trial-averaged smoothed and normalized spike trains and projected the time-varying activity of neurons onto a low-dimensional space. The two principal components that captured the dynamics of putative PYRs (1^st^ component) and INs (2^nd^ component), were strikingly similar across age groups (Figure 4E). This indicates that, while the inhibition strength exerted by INs increases throughout development, the dynamics with which the PYR-IN network responds to IN activation does not change across the first two postnatal weeks. These data provides support to the notion that, on a network level, GABA exerts an inhibitory effect already in the first postnatal week^62,63^.

To corroborate these results, we carried out analogous experiments in mice expressing an inhibitory opsin in INs (Figure 5A, Supp. Figure 3E). Optogenetic inhibition of INs led to conspicuous effects on LFP power, with an increase that was mostly limited to lower frequencies (1-10 Hz) in P2-P6 mice and covered a broader frequency spectrum (1-50 Hz) at older ages (Supp. Figure 3F). Further, inhibiting INs paradoxically tilted E-I ratio towards inhibition and induced an increase in the PSD slope when compared to baseline (main IN inhibition effect, p=0.017, linear model) (Figure 5B). Analogous paradoxical effects induced by IN inhibition have already been reported and are attributed to rebound IN excitation^13^. In support of this hypothesis, the PSD slope computed on the 2^nd^ half of the stimulation tended to increase, yet below significance threshold, when compared to the PSD slope computed on the 1^st^ half of the stimulation (main condition effect, p=0.053, linear mixed-effect model) (Supp. Figure 3G).

**Figure 5.**
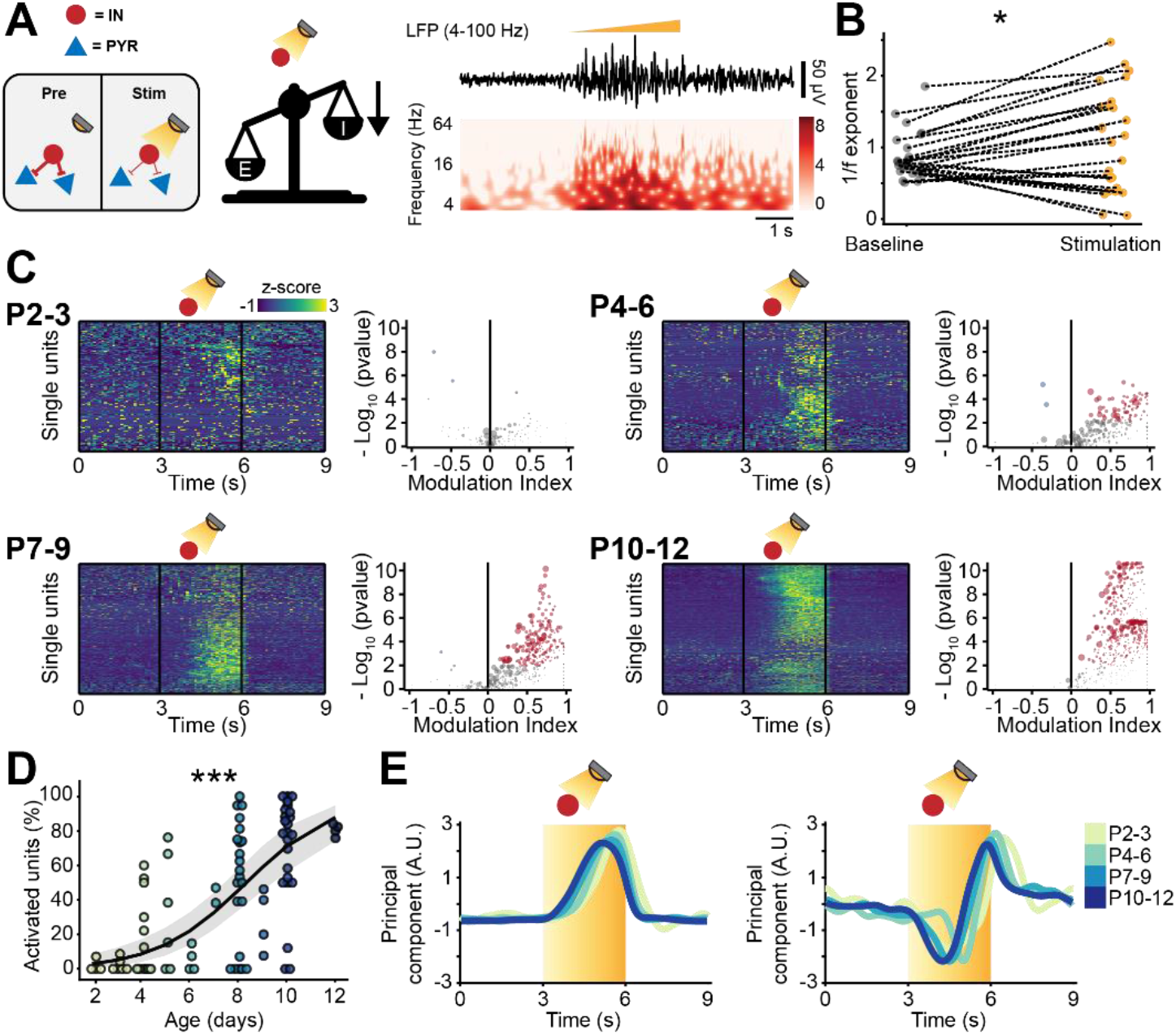
Optogenetic inhibition of IN activity leads to widespread excitation in the developing mouse mPFC. (**A**) Schematic representation of the effects induced by optogenetic IN inhibition (left). Representative LFP trace (4-100 Hz band-pass filter) with a corresponding wavelet spectrum at an identical timescale during ramp light stimulation (594 nm, 3 s) of INs in the mPFC of a P10 mouse. (**B**) Scatter plot displaying the 1/f exponent during baseline and IN optogenetic inhibition. (**C**) Z-scored single unit firing rates in response to optogenetic stimulation of INs (left) and volcano plot displaying the modulation index of pre vs stim single unit firing rates (right) for P2-3 (top left, n=164 units and 5 mice), P4-6 (top right, n=286 units and 11 mice), P7-9 (bottom left, n=470 units and 13 mice) and P10-12 (bottom right, n=691 units and 11 mice) mice. Color codes for firing rate. (**D**) Scatter plot displaying the percentage of activated units with respect to age (n=40 mice). (**E**) 1^st^ (top) and 2^nd^ (bottom) PCA component of trial-averaged spike trains in response to optogenetic inhibition of INs. Color codes for age. In (D) the regression is presented as mean and 95% C.I. Asterisks in (B) and (D) indicate significant effect of IN inhibition and age, respectively. * p < 0.05, *** p < 0.001. Linear model (B) and generalized linear mixed-effect model (D). For detailed statistical results, see S1 Table.

In line with this interpretation, light-induced IN silencing resulted in a widespread progressive increase of firing, with very few inhibited neurons (12/1611 neurons) (Figure 5C). Independent of age group, the increased firing abruptly returned to baseline levels upon terminating the optogenetic stimulus (Figure 5C). The proportion of neurons responding with a firing rate increase during IN inhibition augmented with age (main age effect=0.54, 95% C.I. [0.37; 0.74], p<10^-8^, generalized linear model) (Figure 5D). To qualitatively compare the “neuronal trajectories”, we performed trial-averaged PCA. The 1^st^ component, that captured the activity of putative PYRs, responded to the light stimulus with a monotonic rise of firing rate that quickly dropped as soon as the stimulus stopped (Figure 5E). On the contrary, and in line with previous experimental^13,64,65^ and theoretical work^66,67^ in the adult brain, putative INs (the trajectory captured by the 2^nd^ component), had a biphasic response. They were initially inhibited but, halfway through the optogenetic stimulation, displayed a steep increase in firing rate, that persisted until end of the stimulus (Figure 5E). This “paradoxical” increase in interneuron firing rate emerged more rapidly as mice developed (Figure 5E) and might be the cause of the 1/f exponent shift towards higher values. Thus, while the network excitation derived from IN inhibition increased throughout development, the dynamics with which the PYR-IN network responds to IN inhibition did not change across the first two postnatal weeks. These data provides support to the notion that, on a network level, GABA exerts an inhibitory effect already in the first postnatal week^62,63^.

We have previously shown that the optogenetic paradigm that we utilized does not lead to significant tissue heating^57^, but to further rule out possible nonspecific effects, we applied the same stimulation paradigm to cre-mice (n=10 mice, 380 neurons, Supp. Figure 4A). Pooling together all investigated mice, only 6 out of 380 units were activated, whereas none was inhibited. These results are in line with the statistical threshold (0.01) that was used for this analysis (proportion of modulated units 0.016, C.I. [0.007; 0.034], Supp. Figure 4B). Thus, the used light stimulation leads to minimal, if any, unspecific modulation of neuronal firing.

Taken together, these data show that optogenetic manipulation of INs robustly affects the neonatal prefrontal network in an age-dependent manner. Stimulating INs induced widespread inhibition of putative PYRs, whereas the contrary was true after IN inhibition. Both effects augmented with age. However, the ability of INs to control the cortical inhibition did not qualitatively change during the first two postnatal weeks, resembling adult patterns. These data provide evidence against the long-standing hypothesis of network-level excitatory effects of GABA in the developing mouse cortex.

### Optogenetic manipulation of IN activity impacts pairwise spike train correlations

To investigate the relationship between age-dependent dynamics of inhibition and decorrelation of spike trains, we compared STTC before IN optogenetic manipulation (STTCpre) to STTC during optogenetic manipulation (STTCstim). Considering that STTCpre and STTCstim could only be computed in 3 seconds epochs (times the number of trials), we firstly verified whether STTCpre was a good predictor of “baseline” STTC. Pooling across mice and different IN manipulations, STTCpre robustly correlated with baseline STTC across every investigated lag, from 2.5 ms to 1 s (0.66, [0.48; 0.72] median and min-max Pearson correlation; 0.68 [0.40; 0.71] median and min-max Spearman correlation) (Supp. Figure 5A-B). Further, STTCstim exhibited lower correlation values with baseline STTC across all lags, a first hint that optogenetic IN manipulation affected STTC (Supp. Figure 5A-B).

As predicted by the experimental and modeling results, optogenetic modulation of IN activity affected the STTC values across all investigated timescales (Figure 6A-B). IN stimulation resulted in decreased STTC values (main IN stimulation effect, p<10^-69^, 1 s lag, linear mixed-effect model) (Figure 6C). On the other hand, IN inhibition, even though it might have only transiently (in the first part of the stimulation) increased E-I ratio, it increased STTC (main IN inhibition effect, p<10^-273^, 1 s lag, linear mixed-effect model) (Figure 6D). Moreover, in line with the strongest decorrelation along development for nearby neurons (Figure 3D), IN modulation had a larger impact on STTC values of nearby neurons when compared to pairs that are further apart (IN stimulation*distance interaction, p<10^-4^; IN inhibition*distance interaction p<10^-8^, 1 s lag, linear mixed-effect model) (Figure 6E). Conversely, when fitting a statistical model that included the “spatial configuration” of the neuron pair, we found that it only weakly affected STTC (IN stimulation/inhibition*spatial configuration interaction, p=0.21 and p<0.002, 1 s lag, respectively, linear mixed-effect model) (Supp. Figure 5C-F). Further, when fitting a statistical model that included age as a covariate, we found that age interacted with IN inhibition/stimulation, leading to larger effects of IN manipulation as age increases (IN stimulation*age interaction, p<10^-3^; IN inhibition*age interaction p<10^-24^, 1 s lag, respectively, linear mixed-effect model) (Supp. Figure 5G).

**Figure 6.**
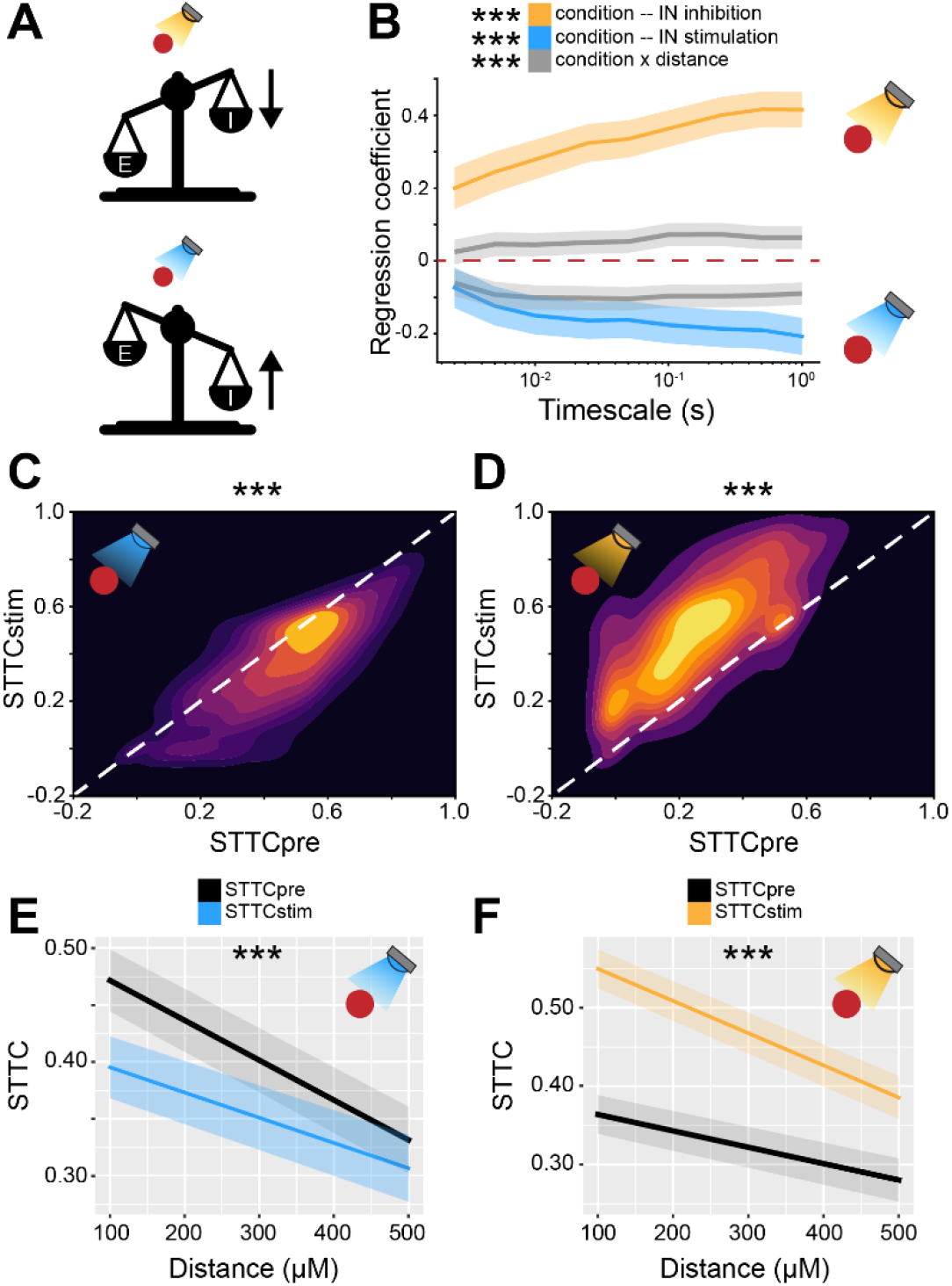
Bidirectional optogenetic manipulation of IN activity affects STTC in the developing mouse mPFC. (**A**) Schematic representation of the effects induced by optogenetic IN inhibition (top) and stimulation (bottom). (**B**) Multivariate linear regression coefficients as a function of STTC lag (n=19951 spike train pairs and 59 mice). (**C** and **D**) 2D kernel density plots displaying STTCpre and STTCstim during IN activation (C) and inhibition (D) (n=10173 spike train pairs and 19 mice, n=9778 spike train pairs and 40 mice, respectively). (**E** and **F**) Average STTCpre and STTCstim during IN activation (E) and inhibition (F) over distance (n=10173 spike train pairs and 19 mice, n=9778 spike train pairs and 40 mice, respectively). In (B) regression coefficients are presented as mean and 95% C.I. In (E and F) data are presented as mean ± SEM. Asterisks in (B) indicate significant regression coefficients of the respective interactions between variables for STTC at 1 s lag Asterisks in (C and D) indicate significant effect of IN activation and inhibition, respectively. Asterisks in (E and F) indicate significant effect of IN activation*distance and IN inhibition*distance interaction, respectively. *** p < 0.001. Linear mixed-effect models. For detailed statistical results, see S1 Table.

Thus, these data indicate that IN manipulation causally impacts pairwise correlations between spike trains. The effect of IN manipulation increases with age, in agreement with the notion that inhibition strengthens throughout development.

### Mouse models with altered developmental E-I ratio have excessively decorrelated activity

Developmental imbalances in E-I ratio have been linked to the pathophysiology of neurodevelopmental disorders^25,58^. A corollary of the results above is that impaired developmental E-I ratio would predict altered correlation levels of neuronal activity. To test this hypothesis, we interrogated two open-source datasets that we recently published^56,58^. They consist of recordings from the prelimbic subdivision of the mPFC from two different developmental mouse models of disease with reduced E-I ratio^56,58^.

The first dataset was obtained from extracellular recordings of SUA from the mPFC of P4-10 control and dual-hit genetic-environmental (GE) mice. GE mice mimic the etiology (combined disruption of *Disc1* gene and maternal immune activation) and cognitive impairment of schizophrenia, showing already at neonatal age reduced excitatory activity in the superficial layers of the mPFC^56,68^ (Figure 7A). Therefore, we hypothesized that GE mice have lower STTC values than controls (i.e. mice lacking the abnormal genetic background and influence of environmental stressor). Considering the layer-specificity of the deficits identified in the mPFC of GE mice, we reasoned that this effect should be present in spike trains from neurons in the superficial layers. Overall, GE mice had lower spike train correlations when compared to controls (main condition effect, p=0.032, 1 s lag, linear mixed-effect model) (Supp. Figure 6A). In line with the proposed hypothesis, this deficit depended on whether the neuron pair was situated in the superficial or deep layers of the mPFC (condition*layer interaction, p < 10^-7^, 1 s lag, linear mixed-effect model). While there was no significant difference between STTC of controls and GE spike train pairs situated in the deep layers (p=0.15, 1 s lag, linear mixed-effect model), spike train pairs of GE mice in which one of the two neurons was located in the superficial layers had reduced STTC values (p=0.016, 1 s lag). This difference was even more robust if both neurons were situated in the superficial layers (p=10^-3^, 1 s lag) (Figure 7B-E). Lastly, the effect did not depend on the age of the mouse (condition*age interaction, p=0.16, 1s lag, linear mixed effect model) (Supp. Figure 6A-B).

**Figure 7.**
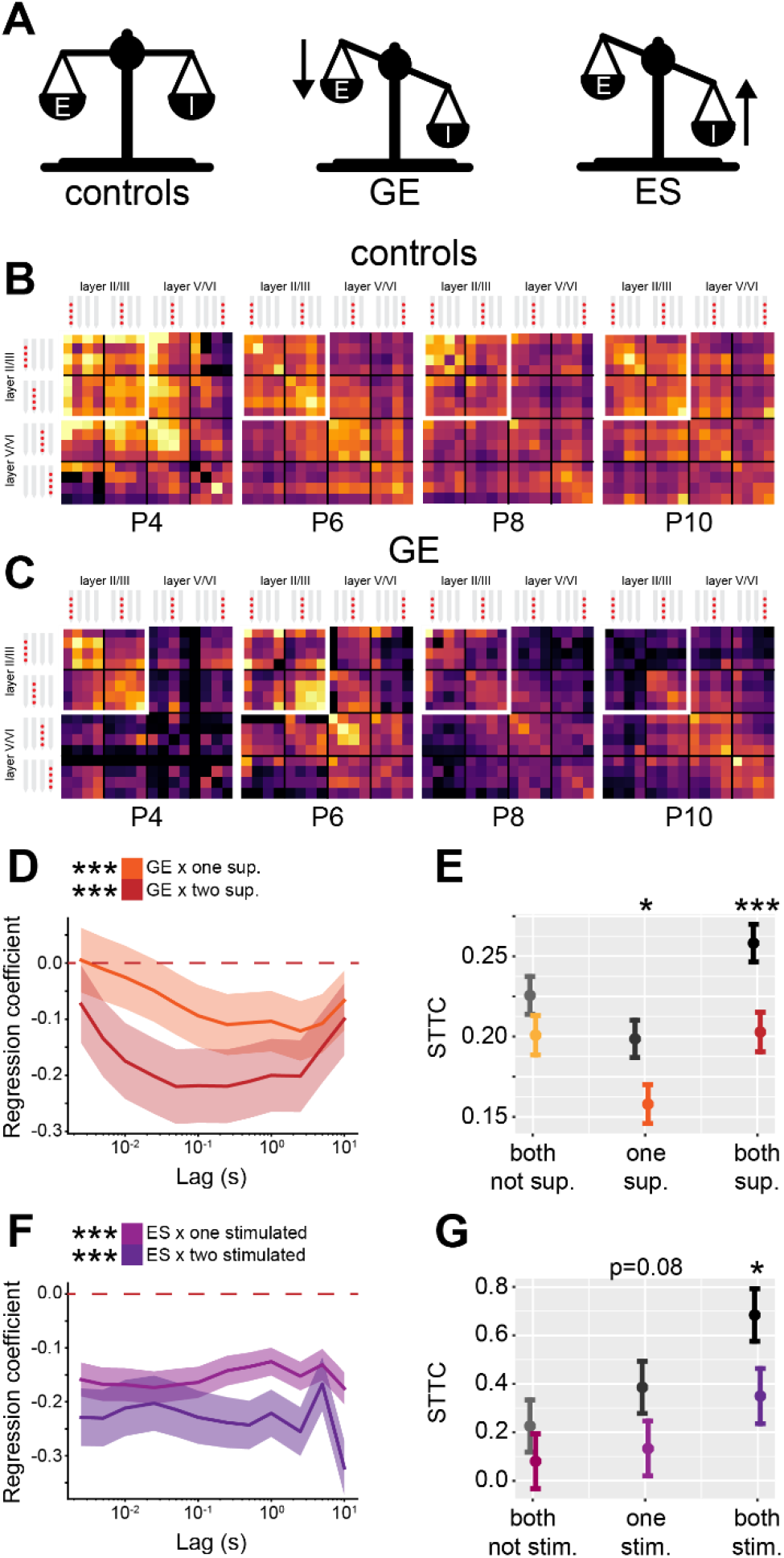
GE and ES mice have reduced STTC values with specific spatial profiles. (**A**) Schematic representation of the E-I ratio imbalance affecting GE and ES mice. (**B**) Weighted adjacency matrices displaying average STTC at 1 s lag of P4, P6, P8, P10 and control mice as a function of the recording sites in which the spike train pair has been recorded (n=18839 spike train pairs and 33 mice). White inset indicates STTC values between spike trains that are located in the superficial layers of the mPFC. Color codes for STTC value. (**C**) Same as (B) for GE mice (n=11051 spike train pairs and 30 mice). (**D**) Multivariate linear regression coefficients as a function of STTC lag (n=29890 spike train pairs and 63 mice). (**E**) STTC of control and GE mice (n=18839 and 11051 spike train pairs; 33 and 30 mice, respectively) with respect to the number of neurons in the superficial layers in the mPFC. (**F** and **G**) Same as (D) and (E) for control and ES mice (n=17150 and 11449 spike train pairs; 10 and 11 mice, respectively) with respect to the number of chronically stimulated neurons. In (D) and (F) regression coefficients are presented as mean and 95% C.I. In (E) and (G) data are presented as mean ± SEM. Asterisks in (D) and (F) indicate significant regression coefficients of the respective interactions between variables for STTC at 1 s lag. Asterisks in (E) and (G) indicate significant difference to control mice. Asterisks in (E and G) indicate significant effect of the respective variable for STTC at 1 s lag. * p < 0.05, *** p < 0.001. Linear mixed-effect models. For detailed statistical results, see S1 Table.

The second dataset originated from extracellular recordings of SUA in the mPFC of P11-12 control and early stimulated (ES) mice. In the mPFC of ES mice a subpopulation of layer II/III PYRs has been transiently stimulated by light during a defined (P7-11) developmental period^58^. We have previously shown that this manipulation causes a long-lasting E-I imbalance and an increased feedback inhibition onto neuronal connections from chronically stimulated PYRs^58^ (Figure 7A). Here we hypothesize that spike trains pairs that involve chronically stimulated PYRs in ES mice should have lower STTC than controls. Overall, ES mice did not show lower levels of STTC (main condition effect, p=0.13, 1 s lag, linear mixed-effect model). However, a strong interaction between condition and the number of neurons in the spike train pair that had been chronically stimulated (condition*number of tagged neurons interaction, p<10^-45^, 1s lag, linear mixed-effect model) was detected. Pairs of spike trains that consisted of two neurons that were not chronically stimulated did not have reduced STTC levels (p=0.36, 1 s lag, linear mixed effect model). However, when one neuron in the pair had been chronically stimulated, the STTC tended to decrease (p=0.078, 1 s lag, linear mixed effect model), the effect becoming more robust when both neurons had been chronically stimulated (p=0.026, 1 s lag, linear mixed effect model) (Figure 7F-G).

Taken together, these data support the hypothesis that decreased developmental E-I ratio results in reduced spike train pairwise correlations. Further, we show that this effect is remarkably specific. In GE mice, a mouse model characterized by reduced excitatory drive in prefrontal PYRs of the superficial layers, the reduced correlation levels were largely limited to spike train pairs involving PYRs of the superficial layers. Similarly, in ES mice, a mouse model characterized by increased feedback inhibition onto chronically stimulated PYRs, the decreased correlation levels were largely limited to spike train pairs involving chronically stimulated PYRs.

### E-I ratio decreases with age in newborn babies

Taking into account the role of E-I ratio for neurodevelopmental disorders, it is of critical relevance to assess whether a developmental strengthening of inhibition occurs also in newborn babies. To this aim, we interrogated two EEG datasets recorded in newborn babies of an age between 35 and 46 post-conception weeks (PCW), a stage of brain development that is roughly equivalent to the one that we studied in mice^1^. While it is not straightforward to compare intracranial recordings from a deep structure like the mouse mPFC to human EEG data, to maximize the consistency between approaches, we limited our analysis to channels from the frontal derivations of the EEG (Figure 8A).

**Figure 8.**
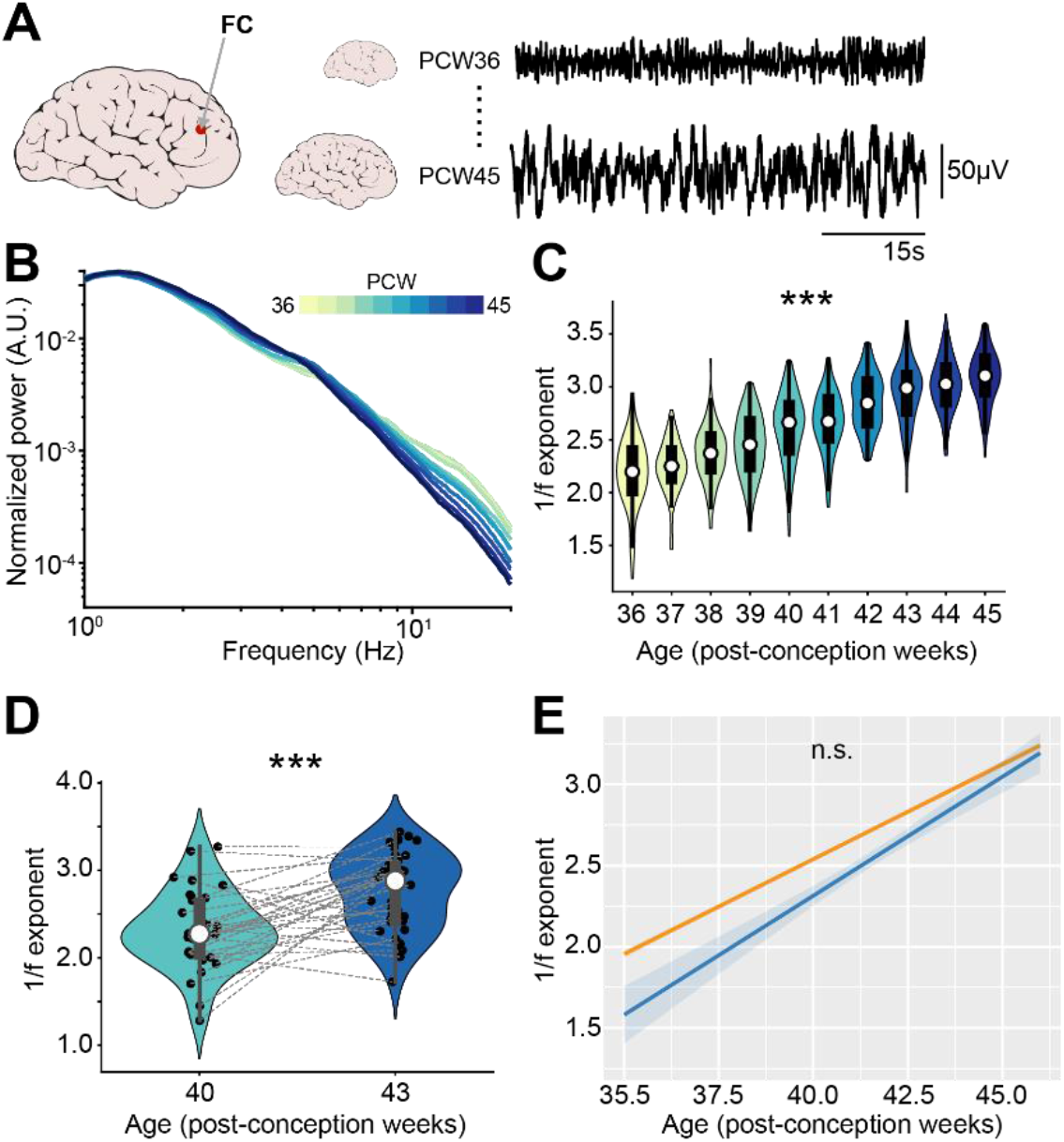
1/f exponent of EEG recordings increases with age in newborn babies. (**A**) Schematic representation of EEG recording from frontal derivations of PCW36-45 newborn babies (left) displayed together with representative EEG traces from PCW36 and PCW45 newborn babies (right). (**B**) Log-log plot displaying the normalized mean PSD power in the 1-20 Hz frequency range of PCW36-45 newborn babies (n=1110 babies). Color codes for age. (**C**) Violin plots displaying the 1/f exponent of PCW36-45 newborn babies (n=1110 babies). (**D**) Same as (C) for PCW40 and PCW43 newborn babies (n=72 EEG recordings and 40 babies). (**E**) 1/f exponent over age for the two EEG datasets (n=1110 babies and n=72 EEG recordings and 40 babies, respectively). In (D) black dots indicate individual data points. In (C) and (D) data are presented as median, 25^th^, 75^th^ percentile and interquartile range. In (C) and (D) the shaded area represents the probability distribution density of the variable. In (B) and (E) data are presented as mean ± SEM. Asterisks in (C) and (D) indicate significant effect of age. *** p < 0.001. Linear model (C) and linear mixed-effect models (D-E). For detailed statistical results, see S1 Table.

The first dataset^69,70^ consisted of 1100 EEG recordings from sleeping babies with an age comprised between 36 and 45 post-conception weeks. Similar to the PSDs of recordings from the neonatal mice mPFC, the PSD slope grew steeper over age (Supp. Figure 7A), a phenomenon that was readily apparent after normalization of the PSD (Figure 8B). We quantified the 1/f exponent and confirmed that it increased with age (age coefficient=0.26, 95% C.I. [0.24; 0.27], p<10^-183^, linear model) (Figure 8C). A second dataset^71^ consisted of EEG recordings from 42 sleeping babies, recorded at 40 and 43 post-conception weeks. The analyses revealed that also for these data the PSD slope grew steeper (Supp. Figure 7B-C) and the 1/f exponent increased with age (age coefficient=0.30, 95% C.I. [0.17; 0.42], p<10^-4^, linear mixed-effect model) (Figure 8D). The increase in 1/f slope over age was very similar across the two different datasets (mean age coefficients=0.26 and 0.30) and no statistical difference was found between them (main dataset effect, p=0.15; age*dataset interaction, p=0.21, linear mixed effect model) (Figure 8E).

Thus, the E-I ratio decreases along development also in newborn humans and might represent a fingerprint of cortical circuit maturation in mammalian species.

## Discussion

Integration of INs into the cortical circuitry has been proposed to bear important structural, functional and behavioral consequences^33^. Here, we show that, even though INs inhibit neuronal activity already in the first postnatal week, the relative strength of the exerted inhibition increases with age, leading to a decrease of E-I ratio. This developmental process is responsible for a transition in brain dynamics, from early highly synchronous activity patterns to decorrelated neural activity later in life. We further show that an early imbalance in E-I ratio in the mPFC results in altered temporal neural dynamics. Lastly, leveraging two EEG datasets recorded in newborn babies, we provide evidence for analogous developmental processes taking place in the murine and human cortex.

Inhibitory synaptogenesis is an exquisitely specific process that is orchestrated by distinct molecular programs^72^ and neural activity^73^. Its timeline is protracted and it extends to early adulthood in the mouse somatosensory cortex^38^. PYRs are thought of playing an instructive role in this process, by providing molecular cues guiding IN migration and by regulating their survival in an activity-dependent manner^33,74^. Early inhibitory circuits have several peculiarities, including a predominance of inhibitory synapses by SOM^+^ INs^35–37^ and the presence of hub neurons that control the formation of functional assemblies^75,76^. One of the unique traits of early inhibitory circuits that has garnered much attention is the hypothesis that GABA might act as an excitatory neurotransmitter. This excitatory action of GABA has been most intensively investigated at single-cell level^77,78^ and has been proposed to result from the low expression of KCC2, a potassium-chloride cotransporter extruding chloride^78^, which leads to high intracellular chloride concentration in immature neurons. The “GABA-switch” from an excitatory to inhibitory neurotransmitter has been suggested to take place between the first and the second postnatal week^78^. Refuting this hypothesis, recent studies have shown that, already during the first postnatal week, GABAergic transmission exerts an inhibitory action on population activity^39,62,63^. In line with this latter interpretation, this study provides SUA-level evidence of the inhibitory effect of GABA *in vivo* already in the first postnatal days. This is of particular interest considering that the PFC is a brain area whose development is thought of being more protracted than sensory areas^1^, where the early effects of GABA have been more intensively studied^39,62,63^. We show that optogenetic inhibition of cortical INs results in a widespread increase of SUA firing rates and, conversely, increasing their activity leads to a reduction of neuronal activity. While the strength of the inhibition exerted by INs increases throughout development, the ability of INs to control cortical inhibition does not qualitatively change with age. Already during the first postnatal week, inhibition of INs leads to a paradoxical increase in their firing rate, This dynamics is characteristic for inhibition-stabilized networks that have a high degree of recurrent inhibition^65^. Thus, while the effects of GABA might differ within brain regions^63^, our data supports the notion that, in the mPFC, GABA has an inhibitory role already at P2.

As the brain develops and inhibitory synaptogenesis progresses, the temporal coordination between excitatory and inhibitory transmission tightens^3,13^ with a relative strengthening of inhibition with respect to excitation (i.e. a decrease in E-I ratio)^12^. While E-I ratio is thought of being a hallmark of neural networks with important implications^13^, the functional consequences of the developmental E-I ratio decrease are still poorly understood. Individually, E-I ratio imbalances^24–29^ and altered correlation structure of brain activity^30–32^ have both been linked to mental disorders. Studying how these two processes are linked to each other is therefore likely to be insightful for understanding the pathophysiology of these disorders. To address this knowledge gap, we explored the impact of varying E-I ratio in a LIF neural network model. In line with previous results^24,46^, we show that E-I ratio can be indirectly tracked by measuring the 1/f exponent, a notion that we leveraged to show that there is an E-I ratio decrease in the mouse and human mPFC. We further show that, in a neural network model, a relative increase in inhibition results in decreased pairwise correlations of spike trains. Confirming the modeling results, we report that, in the mouse mPFC, the E-I ratio decrease taking place across the first two postnatal weeks is accompanied by a reduction in pairwise spike-trains correlations. To further strengthen the link between the two processes, we bidirectionally manipulated the activity of prefrontal INs by light. In line with our hypothesis, optogenetic stimulation of INs (i.e. decreasing E-I ratio) results in reduced correlations among spike trains. Conversely, optogenetic inhibition of INs (i.e. increasing E-I ratio) causes increased correlations among spike trains. IN inhibition results in increased spike-train correlations even though, in the last portion of the optogenetic stimulation, IN display a paradoxical increase in firing rate. This might indicate that even a transient reduction in inhibition strength might be sufficient to increase neural correlations. Both the age-dependent as well as IN manipulation-induced effects on activity correlations do not impact all spike train pairs in a uniform manner. Rather, neuron pairs that are close to each other are more severely affected than those that are farther apart. While an IN subtype-specific dissection of the mechanisms accounting for neural activity decorrelation is beyond the scope of this study, this effect might be explained by the fact that PV^+^ interneurons preferentially provide local inhibition^79^ but have a particularly protracted integration into the prefrontal cortical circuitry^11^. PV^+^ interneurons generally provide inhibition at the soma and the axon initial segment^79^, a position that is particularly suited to inhibit the spiking output of PYRs and thereby reduce pairwise spiking correlations. The progressive embedment of PV^+^ interneurons in the rodent prefrontal circuitry has also been suggested to be responsible for the increase in the average frequency of the LFP oscillations that are generated by layer 2/3 PYRs^11^. Future studies might shed light on whether this process is related to the decorrelation of brain activity.

Here, we show that the 1/f exponent derived from LFP recorded from the mouse mPFC increases along the first two postnatal weeks. Similarly, the 1/f exponent derived from the frontal derivations of EEG recordeds from human babies increases between the 36^th^ and the 45^th^ post-conception week (from ~2 weeks preterm birth to ~7 weeks after birth, when considering 38 weeks as the average length of human pregnancy^80^). This is the opposite of processes taking place in aging^81^, between childhood and adulthood^82^, and even from the 1^st^ to the 7^th^ month of life^83^. The decline in the 1/f exponent (indicative of increased E-I ratio) occurring between childhood and senility can be explained by the decline of brain GABA levels^84^ and cortical inhibition^85^. This is however unlikely to explain the discrepancy between the current study and the effect reported for babies of 1-7 months of age^83^, an age range that borders the one that we investigated. In age-matched mice, a wave of interneuronal cell death takes place around this age^74^ and might induce an early shift from E-I ratio decrease to E-I ratio increase. Whether a similar process occurs in humans too and might explain the discrepancy between the two studies is still unknown and, due to the ethical and technical limitations of invasive recordings in humans, difficult to address.

Several studies have reported E-I imbalances in the mPFC of mouse models of mental disorders and in patients affected by these diseases^24,26,28,29,31,32^. Following this stream of evidence, we investigated a dataset previously obtained from *in vivo* electrophysiological recordings from the mPFC of a mouse mimicking the etiology of schizophrenia^56,58^, generated by combining two mild stressors, a genetic and an environmental one^56,68^. Their synergistic combination results in severe deficits affecting PYRs that reside in the superficial layers of the mPFC. These neurons display a simplified dendritic arborization, a severe reduction in spine density and firing rate^56,68^. A second data set resulted from *in vivo* electrophysiological recordings from the mouse mPFC that experienced a transient stimulation of PYRs by light during a defined developmental time window^58^. This manipulation results in PYRs having an increased dendritic arborization and spine density when compared to controls^58^. We further showed that, compared to controls, synaptic connections involving chronically stimulated PYRs have a long-term shift of EPSCs to IPSCs ratio towards inhibition^58^. Despite the opposite phenotype affecting PYRs in the superficial layers of GE and ES mice, they display similar behavioral symptoms, and diminished mPFC-dependent cognitive abilities in particular^56,58^. This apparent contradiction can be resolved if one looks at the deficits through the lens of E-I ratio. Both mouse models have similarly reduced E-I ratio in the mPFC superficial layers, that leads to analogously excessively reduced STTC values among spike trains. While in sensory areas correlation might limit information carrying capacity, in associative brain areas, like the mPFC, correlations are thought of improving signal readout, and increased correlations have been linked to improved behavioral performance^86^. This data lends support to the hypothesis that imbalances in E-I ratio might be a possible unifying framework for understanding the circuit dysfunction characterizing neuropsychiatric disorders^26,87,88^. In this perspective, it is relevant that the development of E-I ratio can also be quantified, albeit indirectly, also from EEG recordings of newborn babies. The fact that two different EEG datasets yield similar estimations for the age-dependent changes in the 1/f exponent supports the notion that this parameter might be a robust biomarker of E-I ratio development with potential translational relevance.

## Acknowledgments

We thank Sebastian Bitzenhofer, Johanna Kostka, Jastyn Pöpplau and Lingzhen Song for valuable discussions and feedback on the manuscript, P. Putthoff, A. Marquardt and A. Dahlmann for excellent technical assistance. This work was funded by grants from the European Research Council (ERC-2015-CoG 681577 to I.L.H.-O.), EU-project euSNN (MSCA-ITN-H2020-860563 to I.L.H.-O.) and the German Research Foundation (Ha 4466/10-1, Ha4466/11-1, Ha4466/12-1, SPP 1665, and SFB 936 B5 to I.L.H.-O.) and Landesforschungsförderung Hamburg (LFF76, LFF73 to I.L. H.-O.).

## Author Contributions

M.C. and I.L.H.-O. designed the experiments. M.C. carried out the experiments and analyzed the data. M.C. and T.P. carried out neural network modeling. All authors interpreted the data, wrote, discussed and commented on the manuscript.

## Methods

### Data and code availability

LFP and SUA data that were newly generated for this study are available at the following open-access repository: https://gin.g-node.org/mchini/development_EI_decorrelation.

Code supporting the findings of this study is available at the following open-access repository: https://github.com/mchini/Chini_et_al_EI_decorrelation.

### Experimental models and subject details

All experiments were performed in compliance with the German laws and following the European Community guidelines regarding the research animals use. All experiments were approved by the local ethical committee (G132/12, G17/015, N18/015). Experiments were carried out on C57BL/6J, Dlx5/6-Cre (Tg(dlx5a-cre)1Mekk/J, Jackson Laboratory), Gad2-IRES-Cre (Gad2tm2(cre)Zjh, Jackson Laboratory) and ArchT (Ai40(RCL-ArchT/EGFP)-D, Jackson Laboratory) mice of both sexes. Mice were housed in individual cages on a 12 h light/12 h dark cycle, and were given access to water and food ad libitum. The day of birth was considered P0. To inhibit IN activity, mice from the Dlx5/6-Cre and Gad2-IRES-Cre driver lines were crossed with mice from the ArchT reporter line. To stimulate IN activity, P0-P1 mice from the Dlx5/6-Cre and Gad2-IRES-Cre driver lines were injected in the mPFC with a virus encoding for ChR2 (AAV9-Ef1alpha-DIO-hChR2(ET/TC)-eYFP) as previously described^89^. Details on the data acquisition and experimental setup of open-access datasets that were used in this project have been previously published^56,58,69–71^.

### *In vivo* electrophysiology and optogenetics

#### Surgery

In vivo extracellular recordings were performed from the prelimbic subdivision of the mPFC of non-anesthetized P2-P12 mice. Before starting with the surgical procedure, a local anesthetic was applied on the mice neck muscles (0.5% bupivacain / 1% lidocaine). The procedure was carried out under isoflurane anesthesia (induction: 5%; maintenance: 1-3%, lower for older pups, higher for younger pups). Neck muscles were cut to reduce muscle artifacts. A craniotomy over the mPFC (0.5 mm anterior to bregma, 0.1-0.5 mm lateral to the midline) was performed by first carefully thinning the skull and then removing it with the use of a motorized drill. Mice were head-fixed into a stereotactic frame and kept on a heated (37°) surface throughout the entire recording. (Opto)Electrodes (four-shank, 4×4 recording sites, 100 μm between recording sites, 125 μm shank distance; NeuroNexus, MI, USA) were slowly inserted into the prelimbic cortex, at a depth varying between 1.4 and 2 mm depending on the age of the mouse. A silver wire implanted into the cerebellum was used as ground and external reference. Before signal acquisition, mice were allowed to recover for 30-45 minutes, to maximize the quality and stability of the recording as well as single units yield.

#### Signal acquisition

Extracellular signals were acquired and digitized at a 32 kHz sampling rate after band-pass filtering (0.1-9000 Hz) using an extracellular amplifier (Digital Lynx SX; Neuralynx, Bozeman, MO, USA, Cheetah, Neuralynx, Bozeman, MO, USA).

#### Optogenetic stimulation

Optical stimuli were delivered by an Arduino Uno-controlled (Arduino, Italy) diode laser (Omicron, Austria). The delivered light stimuli varied in wavelength (472 nm or 594 nm) according to the experimental paradigm (IN stimulation and inhibition, respectively). Laser power was titrated before signal acquisition and adjusted to the minimum level that induced the desired neuronal response. Typical light power at the fiber tip was measured in the range of 15-40 mW/mm2. Optogenetic stimulations consisted of ramp-like stimuli of 3 s duration as previously described^56–59^. Ramp stimulations were repeated 30-120 times and carried out on the two outmost lateral shanks of the 4-shank electrodes, corresponding to superficial and deep layers of the mPFC.

#### Histological assessment of electrode position

Epifluorescence images of coronal brain sections were acquired post mortem to reconstruct the position of the recording electrode. Only mice in which the electrodes were placed in the correct position were kept for further analysis.

### Neural network modeling

The architecture of the network was set according to Trakoshis et al. (2020)^24^ and is schematically illustrated in Figure 2A. The network was composed of a total of 400 conductance-based leaky integrate-and-fire units, 80% of which were excitatory (E) (N=320) and 20% were inhibitory (I) (N=80). The units in the network were randomly connected with each other, with a connection probability of 0.2 for each type of synaptic connection. Excitatory (E→E, E→I) and inhibitory (I→I and I→E) synapses were mediated by AMPA and GABA conductances, respectively. All baseline parameter values used in the simulations are listed in Table 1. All simulations were performed using Brain2 for Python3.7^90^.

**Table 1.** Parameters of the leaky integrate-and-fire network.

| Neuron model |  |  |  |
| --- | --- | --- | --- |
| *Parameter* | *Description* | *Excitatory cells* | *Inhibitory cells* |
| $V_L$ | Leak membrane potential | -70 mV | -70 mV |
| $V_{Thr}$ | Spike threshold potential | -52 mV | -52 mV |
| $V_{Res}$ | Reset potential | -59 mV | -59 mV |
| $\tau_{Ref}$ | Refractory period | 2 ms | 1 ms |
| $C_m$ | Membrane capacitance | 500 pF | 500 pF |
| $g_L$ | Membrane leak conductance | 25 nS | 20 nS |
| $\tau_m$ | Membrane time constant | 20 ms | 10 ms |
| Synapse model |  |  |  |
| *Parameter* | *Description* | *Excitatory cells* | *Inhibitory cells* |
| $E_{AMPA}$ | Reversal potential (AMPA) | 0 mV | 0 mV |
| $E_{GABA}$ | Reversal potential (GABA) | -80 mV | -80 mV |
| $g_{AMPA}$ | Conductance (AMPA) | 0.178 * 5 nS | 0.254 * 5 nS |
| $g_{GABA}$ | Conductance (GABA) | 2.01 * nS | 2.7 * 5 nS |
| $g_{AMPA,ext}$ | Conductance external input (AMPA) | 0.234 * 5 nS | - |
| $\tau_{AMPA}$ | Time constant of AMPA decay | 2 ms | 1 ms |
| $\tau_{GABA}$ | Time constant of GABA decay | 8 ms | 8 ms |

The dynamics of each excitatory and inhibitory cell were governed by the following stochastic differential equation:

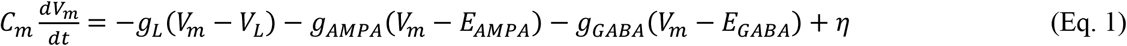

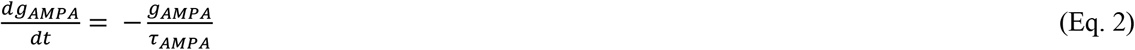

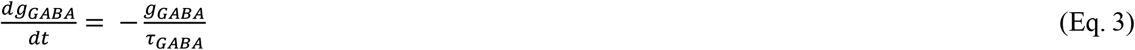

where *V_m_* is the membrane potential, *V_L_* is the leak membrane potential and *E_AMPA_* and *E_GABA_* denote the AMPA and GABA current reversal potentials, respectively. The synaptic conductance parameters and the corresponding decay time constants are denoted by *g_AMPA_, g_GABA_* and *τ_AMPA_, τ_GABA_*, respectively. *η* is a noise term that is generated by an Ornstein-Uhlenbeck process with zero mean. Due to the near-instantaneous rise times of AMPA- and GABA-mediated currents (both typically < 0.5 ms), we opted to neglect these in the current simulations. Moreover, synaptic transmission was assumed to be instantaneous (i.e., with zero delay). The excitatory units of the network received an additional external input in the form of AMPA-mediated Poisson spike trains from an external pool of 100 units with a constant spike rate of 1.5 spikes / second.

In order to assess the effect of altered E-I ratio (*g_E_/g_I_*), we parametrically modulated all excitatory (through multiplication with 21 linearly spaced values from 0.75 to 3) and all inhibitory (21 linearly spaced values from 0.25 to 1.5) synaptic conductances. The network was simulated for a duration of 30 s for each of the 21×21 parameter combinations. For each parameter combination, the LFP of the network was computed by taking the sum of the absolute values of the AMPA and GABA currents on all excitatory cells^24^. Neuronal correlation was estimated by means of the spike time tiling coefficient (see below), assessed at a lag of 1s.

### Electrophysiological analysis

Data were analyzed with custom-written algorithms in the MATLAB and Python environment that are available on the following github repository: https://github.com/mchini/Chini_et_al_EI_decorrelation.

#### Detection of active periods

During early development, brain activity is characterized by an alternation of periods of isoelectric traces (silent periods) and oscillatory bursts (active periods). To detect and quantify the properties of active periods, we developed a novel detection algorithm. For this, the extracellular signal was band-pass filtered (4-20 Hz) and downsampled to 100 Hz, before being averaged across recording electrodes. The average signal (raw and z-scored) was then passed through a boxcar square filter (500ms) on which a hysteresis threshold was applied. Active periods were firstly detected as oscillatory peaks exceeding an absolute or relative threshold (100 μV or 4 standard deviations, respectively) and subsequently extended to all neighboring time points that exceed a lower threshold (50μV or 2 standard deviations, respectively). The combination of absolute and relative thresholding makes this approach suitable to a wide range of signals, from the highly discontinuous brain activity of P2 mice, to the nearly continuous brain activity of P11-12 mice (Figure 1A-B). Neighboring active periods whose inter-active period-interval was shorter than 1s were merged. Active periods whose duration was smaller than 300ms were discarded.

#### Power spectral density (PSD)

PSDs for mouse and human data (see below for exception) were computed with the *mtspecgramc* function of the Chronux Toolbox (10s long-windows, 5s overlap). Median averaging was the preferred measure of central tendency^91^. To quantify the PSD modulation by IN optogenetic stimulation/inhibition, we computed the MI (see below) of the PSD computed on the last 1.5 s of the optogenetic stimulation with the PSD computed on the 1.5s preceding stimulus delivery.

#### EEG preprocessing

EEG signal was extracted only from frontal electrodes (Fp1, F7, F3, Fp2, F8, F4, Fpz, when available) and re-referenced to a common average reference before further analysis. From the EEG dataset of 1100 sleeping babies^69^, epochs whose average envelope amplitude exceeded two standard deviations from the mean were considered as possible artefacts and were removed from further analysis. No preprocessing was applied to the EEG dataset of sleeping babies recorded at 40 and 43 post-conception weeks, as PSDs were already included in the freely available data^71^.

#### 1-f exponent

The 1-f exponent was extracted on the 5-20 Hz and 5-45 Hz (human and mouse data, respectively) frequency range of PSDs using the FOOOF package^47^ with the “fixed” aperiodic mode. To quantify the 1/f exponent modulation by IN optogenetic stimulation/inhibition, we compared the exponent obtained by PSDs computed on the second half of the optogenetic stimulation with the baseline exponent.

#### Spike sorting

Spike sorting was performed using Klusta^92^. Automatically-obtained clusters were then manually curated using phy (https://github.com/cortex-lab/phy).

#### Spike-Time Tiling Coefficient (STTC)

The STTC, a metric that tracks correlations between spike trains and is robust to changes in firing rate, was calculated as previously described^51,93^ (Figure 3A):

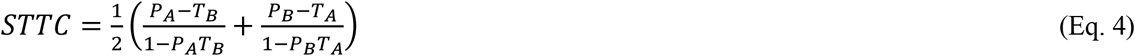

where P_A_ is defined as the proportion of spikes in spike train A that falls within ±Δt of a spike from spike train B. T_A_ is defined as the proportion of time that occurs within (is “tiled” by) ±Δt from the spikes of spike train A. The same applies for P_B_ and T_B_. The “lag” parameter ±Δt controls the “timescale” at which the STTC is computed, a parameter that we systematically varied across more than three orders of magnitude (from 2.5 ms to 10 s). Baseline STTC analysis was limited to spike trains pairs that were recorded for at least an hour and for which both spike trains had at least 50 spikes (40921 of 56613 spike train pairs). To quantify the STTC modulation by IN optogenetic stimulation/inhibition, we compared the STTC derived by spike matrices obtained during the 3s optogenetic stimulation with the STTC derived by spike matrices obtained during the 3s preceding optogenetic stimulation.

#### Spatial arrangements of spike train pairs

To refine our statistical modeling of the sparsification of neural data, we encoded the spatial arrangement of spike train pairs as “local”, “lateral”, “translaminar” or “other” (Figure 3A). Spike trains pairs that consisted of neurons recorded on the same recording site were defined as “local”. Spike trains pairs that consisted of neurons recorded on the same shank (putatively belonging to the same layer) were defined as “lateral”. Spike trains pairs that consisted of neurons recorded on different shanks but at the same depth (putatively belonging to the same “column”) were defined as “translaminar”. Spike train pairs that did not satisfy any of the previous conditions were defined as “other”.

#### Modulation Index (MI)

The MI is a normalization strategy that was used to compute optogenetically-induced changes in firing rate and LFP power. The MI has the desirable property of bounding the normalized value between −1 and 1. MI was computed as:

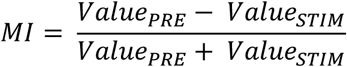

#### Optogenetic modulation of electrophysiological parameters

Modulation of firing rate by optogenetic manipulation was quantified using the MI and signed-rank testing that compared the firing rate during the last 1.5s of optical stimulation with the firing rate during the 1.5s preceding stimulus delivery.

#### PCA of spike matrices during optogenetic stimulations

The first two PCA components of the spike during optogenetic stimulations were computed on trial-averaged spike trains that were convolved with a gaussian window (500 ms length, 50 ms standard deviation) and z-scored across the time dimension.

### Statistical modeling

Statistical modeling was carried out in the R environment. All the scripts and the processed data on which the analysis is based are available from the following github repository: https://github.com/mchini/Chini_et_al_EI_decorrelation.

Nested data were analyzed with (generalized) linear mixed-effects models (*lmer* and *glmer* functions of the *lme4* R package^94^). Depending on the specific experimental design, we used “mouse” or “subject” as random effects. For statistical analysis of STTC, to maximize interpretability of the results (i.e. avoid multiple triple interactions), each STTC lag was investigated as its own independent variable, using identical statistical models. Regression on data that, upon visual inspection seemed to be better fit by an exponential curve, were fitted with generalized linear (mixed-effect) models (family=Gamma, α=1, link=inverse). Proportions (e.g. the proportion of activated/inhibited units) were also fitted with generalized linear (mixed-effect) models (family=Binomial, link=logit). Statistical significance for linear mixed-effects models were computed with the *lmerTest* R package^95^, using the Satterthwaite’s degrees of freedom method. When possible, model selection was performed according to experimental design. When this was not possible, models were compared using the *compare_performance* function of the *performance* R package^96^, and model choice was based on an holistic comparison of AIC, BIC, RMSE and R2. Model output was plotted with the *plot_model* (type=’pred’) function of the *sjPlot* R package^97^. 95% confidence intervals were computed using the *confint* R function. Post hoc analysis with Tukey multiple comparison correction was carried out using the *emmeans* and *emtrends* functions of the *emmeans* R package^98^.

---

**Supp. Figure 1.**
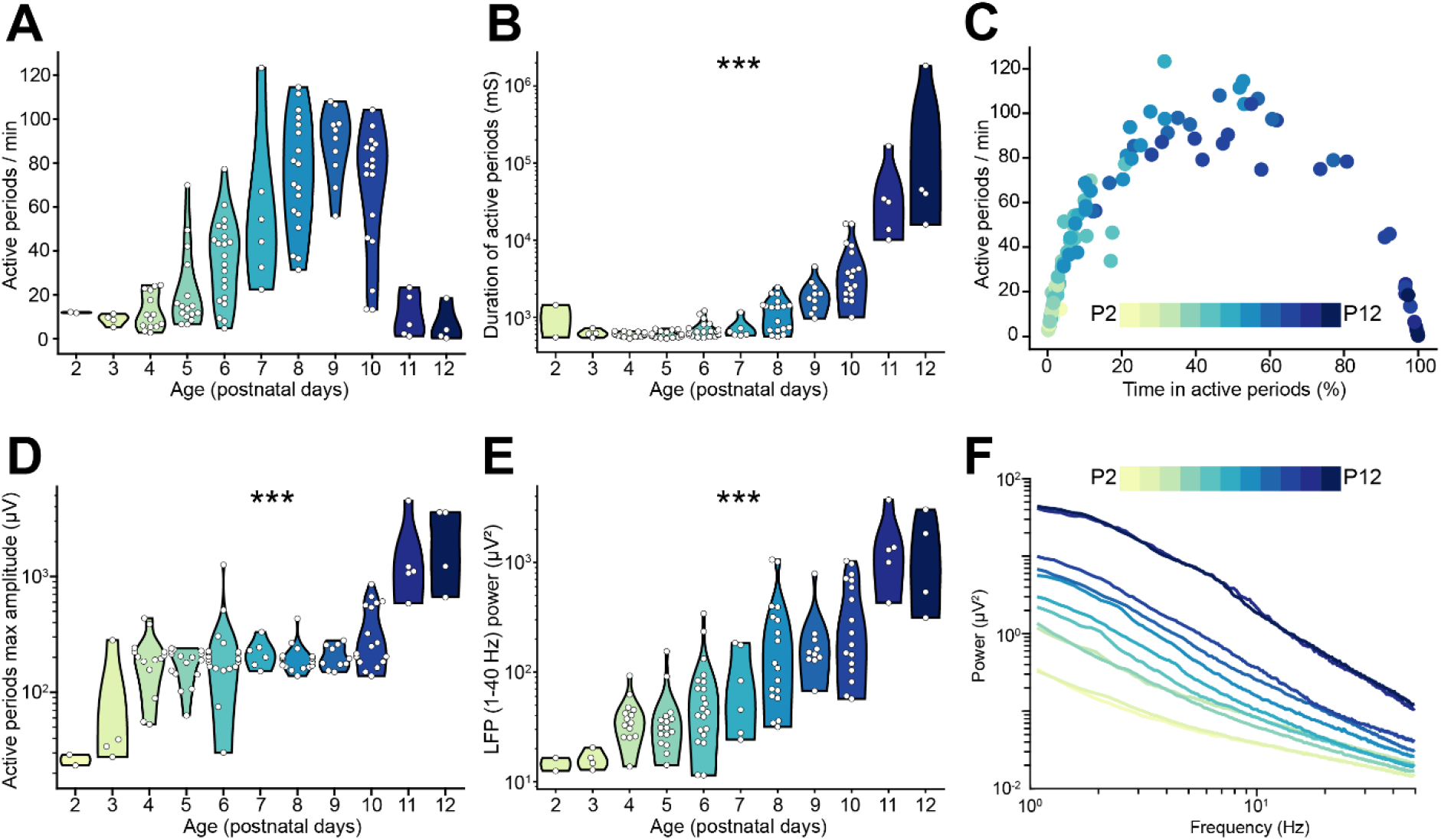
related to Figure 1. LFP and SUA properties of the mouse mPFC across the first two postnatal weeks. (**A** and **B**) Violin plots displaying the number (A) and duration (B) of active periods of P2-12 mice (n=117 mice). (**C**) Scatter plot displaying the time spent in active periods with respect to the number of active periods per minute of P2-12 mice (n=117 mice). Color codes for age. (**D** and **E**) Violin plot displaying the maximum amplitude of active periods (D) and the LFP power in the 1-40 Hz frequency range (E) of P2-12 mice (n=117 mice). (**F**) Log-log line plot displaying the median LFP power in the 0.1-4 Hz frequency range of P2-12 mice (n=117 mice). Color codes for age. In (A), (B), (D) and (E) white dots indicate individual data points and the shaded area represents the probability distribution density of the variable. In (F) data are presented as median. Asterisks in (B), (D) and (E) indicate significant effect of age. *** p < 0.001. Generalized linear models. For detailed statistical results, see S1 Table.

**Supp. Figure 2.**
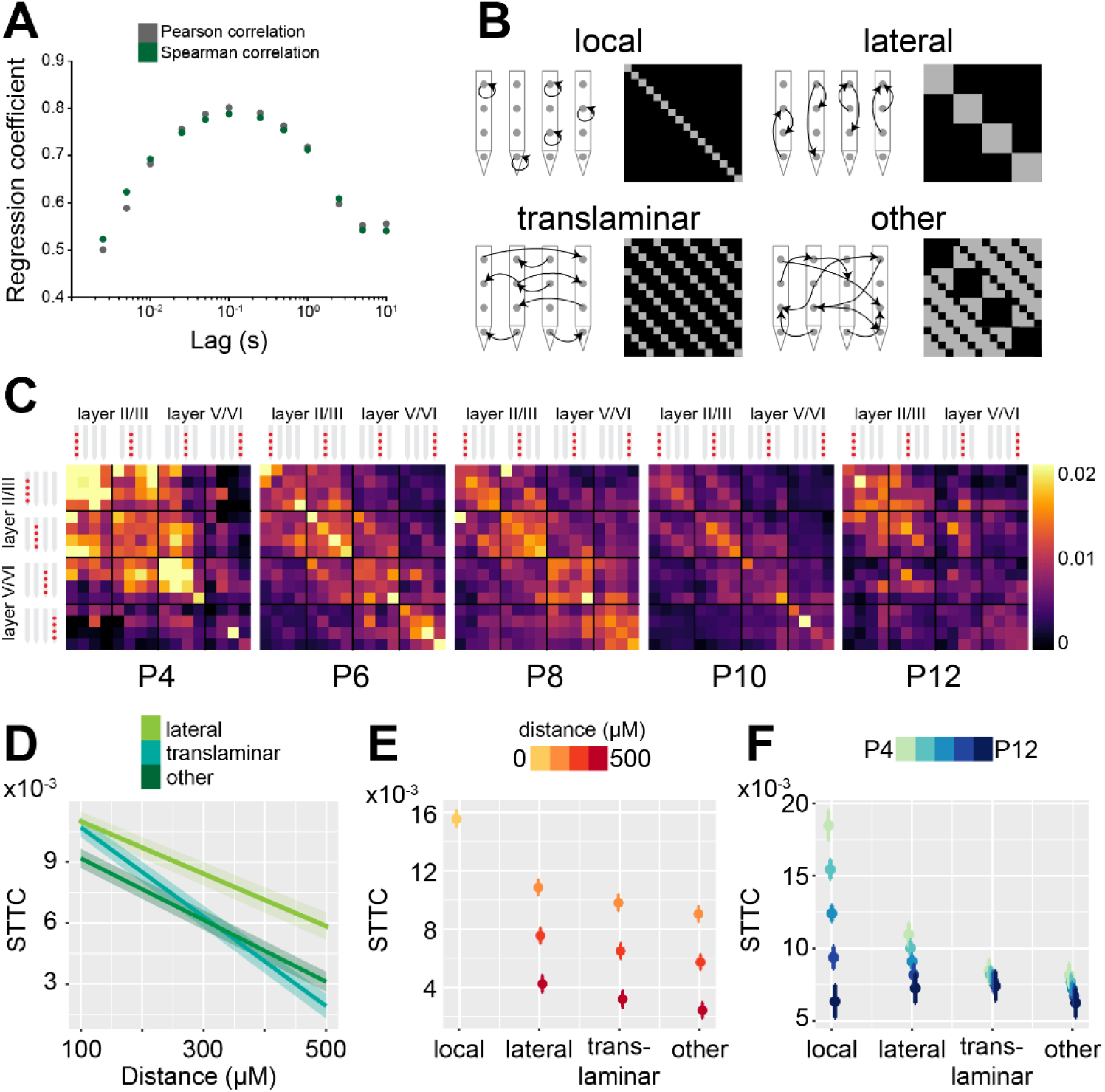
related to Figure 3. The STTC developmental decrease follows specific spatial patterns in the mouse mPFC. (**A**) Scatter plot displaying Pearson and Spearman correlation coefficient for STTC computed on the 1^st^ and 2^nd^ half of the recording over lags (n=40921 spike train pairs and 82 mice). (**B**) Schematic illustration of the spatial configurations of spike train pairs. (**C**) Weighted adjacency matrices displaying average STTC at 2.5 ms lag of P4, P6, P8, P10 and P12 mice as a function of the recording sites in which the spike train pair has been recorded. Color codes for STTC value. (**D**) Average STTC at 2.5 ms over distance (n=40921 spike train pairs and 82 mice). Color codes for spatial configuration. (**E** and **F**) Scatter plot displaying STTC as a function of spatial configuration (n=40921 spike train pairs and 82 mice). Color codes for distance (E) and age (F). In (D-F), data are presented as mean ± SEM. Linear mixed-effect models. For detailed statistical results, see S1 Table.

**Supp. Figure 3.**
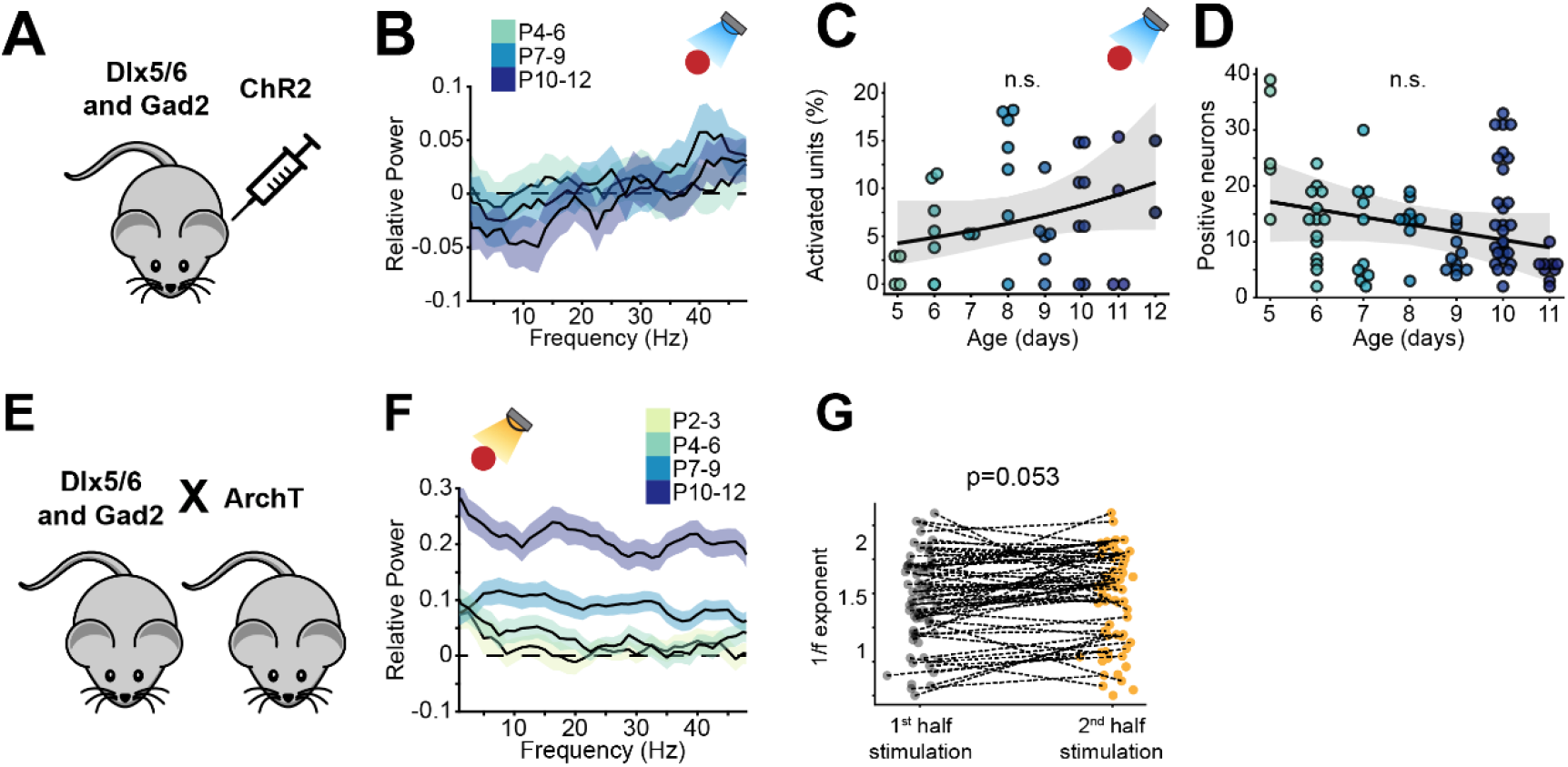
related to Figures 4 and 5. Optogenetic manipulation of IN activity affects LFP activity in the mouse developing mPFC. (**A**) Schematic representation of the experimental approach employed to target INs with an excitatory opsin. (**B**) Relative LFP power in response to ramp light stimulation of INs (n=19 mice). Color codes for age group. (**C**) Proportion of activated units in response to ramp light stimulation of INs as a function of age (n=19 mice). (**D**) Number of virally transfected neurons as a function of age (n=87 images and 16 mice). (**E**) Schematic representation of the experimental approach employed to target INs with an inhibitory opsin. (**F**) Relative LFP power in response to ramp light inhibition of INs (n=40 mice). Color codes for age. (**G**) Scatter plot displaying the 1/f exponent during the 1^st^ and 2^nd^ half of the IN optogenetic inhibition. In (B) and (F) data are presented as mean ± SEM. In (C) and (D) the regression is presented as mean and 95% C.I. Generalized linear mixed-effect model (C) and linear mixed-effect model (D). For detailed statistical results, see S1 Table.

**Supp. Figure 4.**
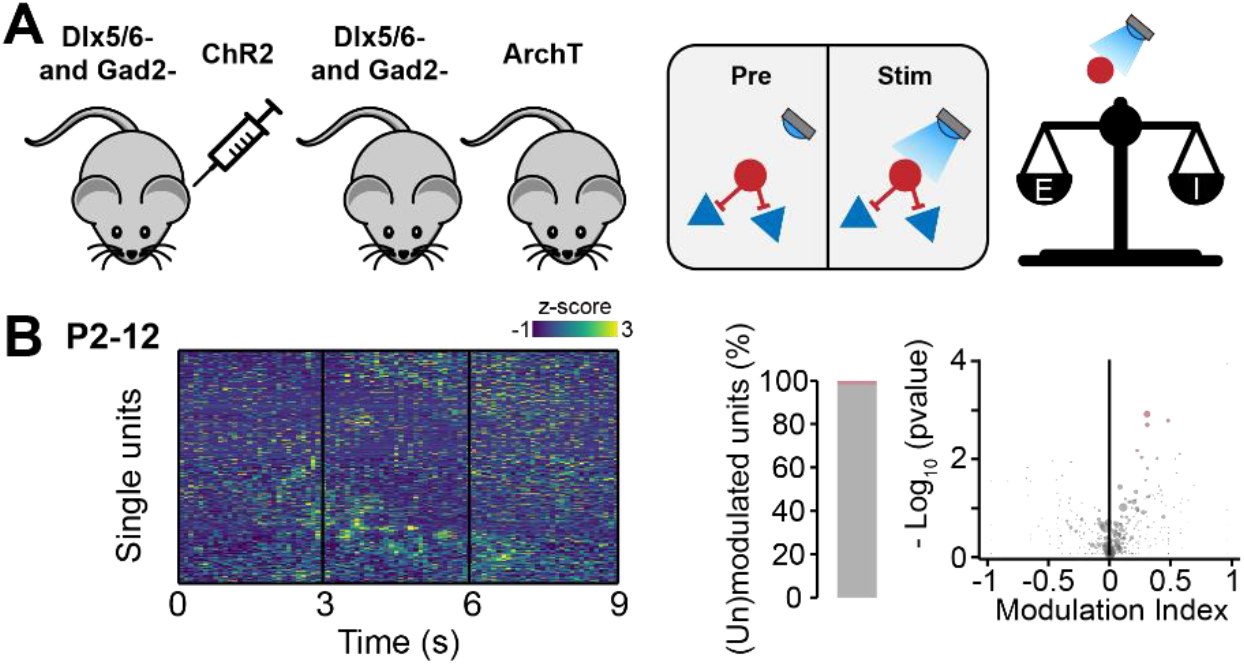
related to Figures 4 and 5. Optogenetic manipulation of IN activity in cre-mice does not affect the developing mouse mPFC. (**A**) Schematic representation of the experimental approach employed to generate control (cre-) mice (left) and schematic representation of the lack of effects induced by optogenetic IN stimulation in cre-mice (right). (**B**) Z-scored single unit firing rates response to optogenetic stimulation in cre-mice (left) and volcano plot displaying the modulation index of pre vs stim single unit firing rates (right) for P2-12 mice (n=380 units and 10 mice),

**Supp. Figure 5.**
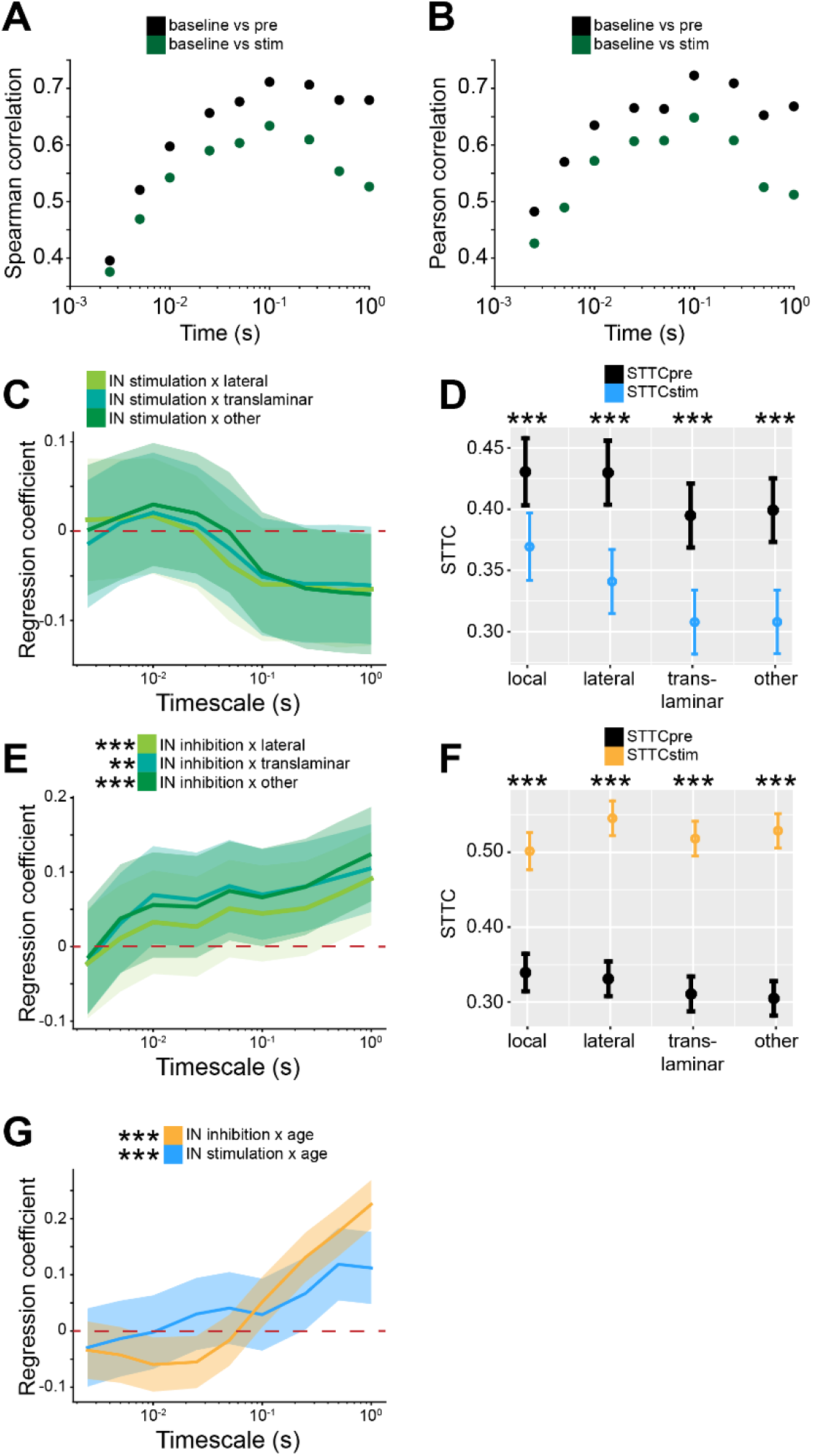
related to Figure 6. Bidirectional optogenetic manipulation of IN activity effects on STTC depends on spatial configurations and age. (**A** and **B**) Scatter plot displaying Pearson (A) and Spearman (B) correlation coefficients for STTCpre and STTCstim with STTC computed on baseline data over lags (n=19951 spike train pairs and 59 mice). (**C**) Multivariate linear regression coefficients for IN stimulation interaction with spatial configuration as a function of STTC lag (n=10173 spike train pairs and 19 mice). (**D**) STTCpre and STTCstim with respect to spatial configuration (n=10173 spike train pairs and 19 mice). (**E** and **F**) Same as (C and D) for IN inhibition (n=9778 spike train pairs and 40 mice). (**G**) Multivariate linear regression coefficients for IN stimulation (blue) and IN inhibition (yellow) interaction with age as a function of STTC lag (n=10173 spike train pairs and 19 mice and n=9778 spike train pairs and 40 mice, respectively). Asterisks in (D and F) indicate significant effect of IN stimulation (D) and inhibition (F). Asterisks in (E and G) indicate significant regression coefficients of the respective interactions between variables for STTC at 1 s lag. ** p < 0.001; *** p < 0.001. In (C, E and G) regression coefficients are presented as mean and 95% C.I. In (D and F) data are presented as mean ± SEM. Linear mixed-effect models. For detailed statistical results, see S1 Table.

**Supp. Figure 6.**
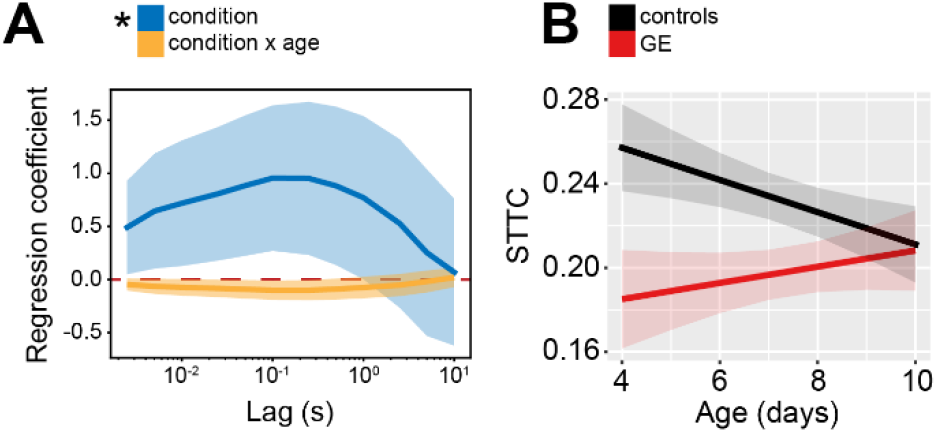
related to Figure 7. Age and spatial profile of STTC values in GE and ES mice. (**A**) Multivariate linear regression coefficients as a function of STTC lag (n=29890 spike train pairs and 63 mice). (**B**) STTC of control and GE mice (n=18839 and 11051 spike train pairs; 33 and 30 mice, respectively) over age. Asterisks in (A) indicate significant regression coefficients of the respective variable for STTC at 1 s lag. * p < 0.05. In (A) regression coefficients are presented as mean and 95% C.I. In (B) and data are presented as mean ± SEM. Linear mixed-effect models. For detailed statistical results, see S1 Table.

**Supp. Figure 7.**
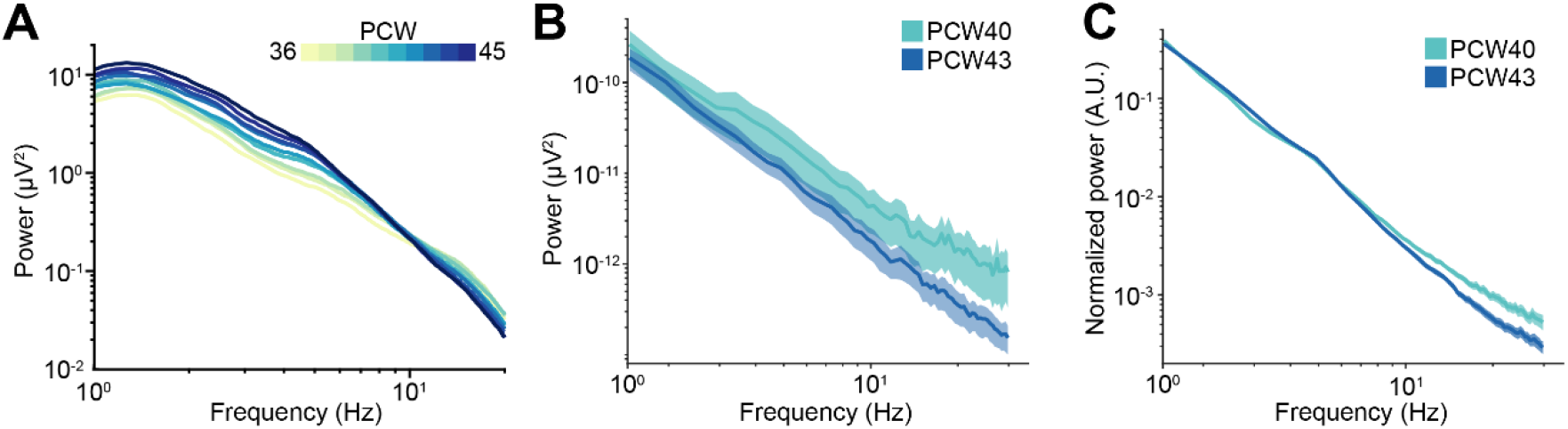
related to Figure 8. EEG PSDs of newborn babies. (**A**) Log-log plot displaying the mean PSD power in the 1-20 Hz frequency range of PCW36-45 newborn babies (n=1110 babies). Color codes for age. (**B**) Same as (A) for PCW40 and PCW43 newborn babies (n=72 EEG recordings and 40 babies). Color codes for age. (**C**) Same as (B) for normalized mean PSD power. In (A-C) data are presented as mean ± SEM.

